# UFMylation anchors splicing factors at the ER to reprogram nuclear splicing

**DOI:** 10.64898/2026.03.30.715226

**Authors:** Ni Zhan, Ranjith K. Papareddy, Bu Erte, Aleksandra S. Anisimova, Catarina Perdigao, Marilyn Tirard-Thevenoud, Milica Mihailovic, Hasan Akyol, G. Elif Karagöz, Nils Brose, Nicholas A.T. Irwin, Yasin Dagdas

**Affiliations:** Gregor Mendel Institute (GMI), Austrian Academy of Sciences, Vienna BioCenter (VBC), Vienna, Austria; Heidelberg University, Centre for Organismal Studies (COS), 69120 Heidelberg, Germany; Max Perutz Labs, Medical University of Vienna, Vienna BioCenter (VBC), Vienna, Austria; Vienna BioCenter PhD Program, Doctoral School of the University of Vienna and Medical University of Vienna, Vienna, Austria; Department of Molecular Neurobiology, Max Planck Institute for Multidisciplinary Sciences, Göttingen, Germany; Cluster of Excellence GreenRobust, Heidelberg University, 69120 Heidelberg, Germany

## Abstract

How organelles communicate stress to the nucleus to coordinate adaptive responses remains a fundamental question in cell biology. Here, we identify a non-canonical retrograde signaling pathway in which stalling-induced UFMylation of ER-associated ribosomes anchors splicing regulators at the ER, directly coupling translational stress to nuclear RNA processing. Phylogenetic profiling linked the UFMylation machinery to a network of nuclear mRNA processing factors. Fractionation-based quantitative proteomics further supported this link and revealed that translational stress triggers UFM1-dependent retention of serine/arginine-rich (SR) splicing factors at the ER, depleting their nuclear pools. Mechanistically, UFMylated ribosomes physically tether SR proteins at the ER surface, driving widespread intron retention that preferentially targets transcripts encoding membrane lipid metabolism and endomembrane-associated processes—a response conserved from plants to mammals. These findings reframe UFMylation from a local ribosome repair signal to a systems-level coordinator of ER-nucleus communication that reprograms nuclear splicing and reshapes membrane-associated gene expression with implications for diverse human diseases linked to UFMylation defects.

## Introduction

The endoplasmic reticulum (ER) is responsible for the synthesis of nearly one-third of the cellular proteome and is the primary site for membrane-bound and secretory protein biogenesis^1−3^. Disruption of ER function perturbs proteostasis; when ribosomes stall or misfolded proteins accumulate, cells activate multiple quality control mechanisms, including the unfolded protein response (UPR), ER-associated degradation (ERAD), selective autophagy of ER fragments (ER-phagy), and ribosome-associated quality control (ER-RQC)^3−7^. Unlike cytosolic RQC^3,8−10^, ribosome stalling at the ER occurs at the SEC61 translocon, where stalled 60S subunits obstruct both the translocon and the ribosomal exit tunnel, creating a unique surveillance challenge. A defining feature of ER-RQC is UFMylation, a ubiquitin-like post-translational modification^11,12^. At the ER, the transmembrane protein DDRGK1 and CDK5RAP3 (C53) serve as adaptors for UFMylation^13^. Upon ribosome stalling, UFL1–C53–DDRGK1 forms an E3 ligase complex that conjugates UFM1 to ribosomal protein RPL26^14−16^, facilitating release of the 60S subunit and enabling LTN1-mediated degrdation of the arrested nascent chain^17−20^. Beyond this core role, UFMylation promotes lysosomal clearance of translocon-arrested proteins via translocation-associated quality control (TAQC) and activates C53-mediated ER-phagy^4,21,22^.

Physiologically, our previous studies demonstrated that the UFMylation pathway is essential for maintaining ER homeostasis in plants, with knockout mutants exhibiting hypersensitivity to ER and salt stress^4,21,23^. In animal cells UFMylation has been implicated in diverse processes, including lipid metabolism^24^, ER–Golgi trafficking^25^, ER expansion^26^, and development^27^, while its disruption leads to embryonic lethality in mice and various human pathologies^11,27,28^. These established roles of suggested that UFMylation coordinate broader cellular adaptations to ER translational stress, beyond the local resolution of ribosome stalling. Indeed, two recent genome-wide CRISPR screens in human iPSC-derived neurons independently identified UFMylation as major modifiers of tau aggregation and propagation^29,30^. While paradoxical at first glance, given that tau is a cytosolic protein, these studies also suggest UFMylation plays a role beyond ER-RQC.

Here, we uncover a mechanism that resolves this paradox. Phylogenetic profiling revealed an unexpected coevolutionary link between UFMylation machinery and nuclear mRNA processing factors. Further mechanistic analyses show that ribosome stalling triggers retention of SR splicing factors at UFMylated ER-bound ribosomes, depleting their nuclear pools. This spatial redistribution rewires the splicing landscape, with intron retention emerging as the predominant outcome across plants, human cells, and mouse neurons. The affected transcripts are enriched for genes encoding membrane lipid metabolism and endomembrane processes, suggesting a feedback circuit for membrane remodeling under stress. Our findings define a retrograde signaling pathway coupling translational surveillance to mRNA splicing.

## Results

### Phylogenetic profiling and nuclear proteomics link UFMylation with nuclear mRNA processing

Previous studies have identified coevolutionary relationships among the canonical components of the UFMylation cascade^21,31^. To examine UFMylation function more broadly, we used phylogenetic profiling to identify additional UFM1-associated proteins. Correlating the presence-absence patterns of UFM1 with eukaryotic protein families across diverse taxa revealed 103 putatively co-evolving protein families (Fig. 1a, b). As expected, these included the core UFMylation machinery (UFL1, UFC1, DDRGK1, C53, UBA5, and UFSP2) and known associated factors such as ODR4^32^ (Fig. 1c). Gene ontology analysis of the co-evolved proteins revealed functional enrichments beyond ER homeostasis, including actin related Scar-WAVE complex, double strand break repair (the Fanconi anemia complex), cytokinesis, and RNA processing (Fig. 1c and Fig. S1a, Table S1). The link to DNA damage response is consistent with recent studies linking UFMylation to genome stability and DNA repair^11^.

**Figure 1.**
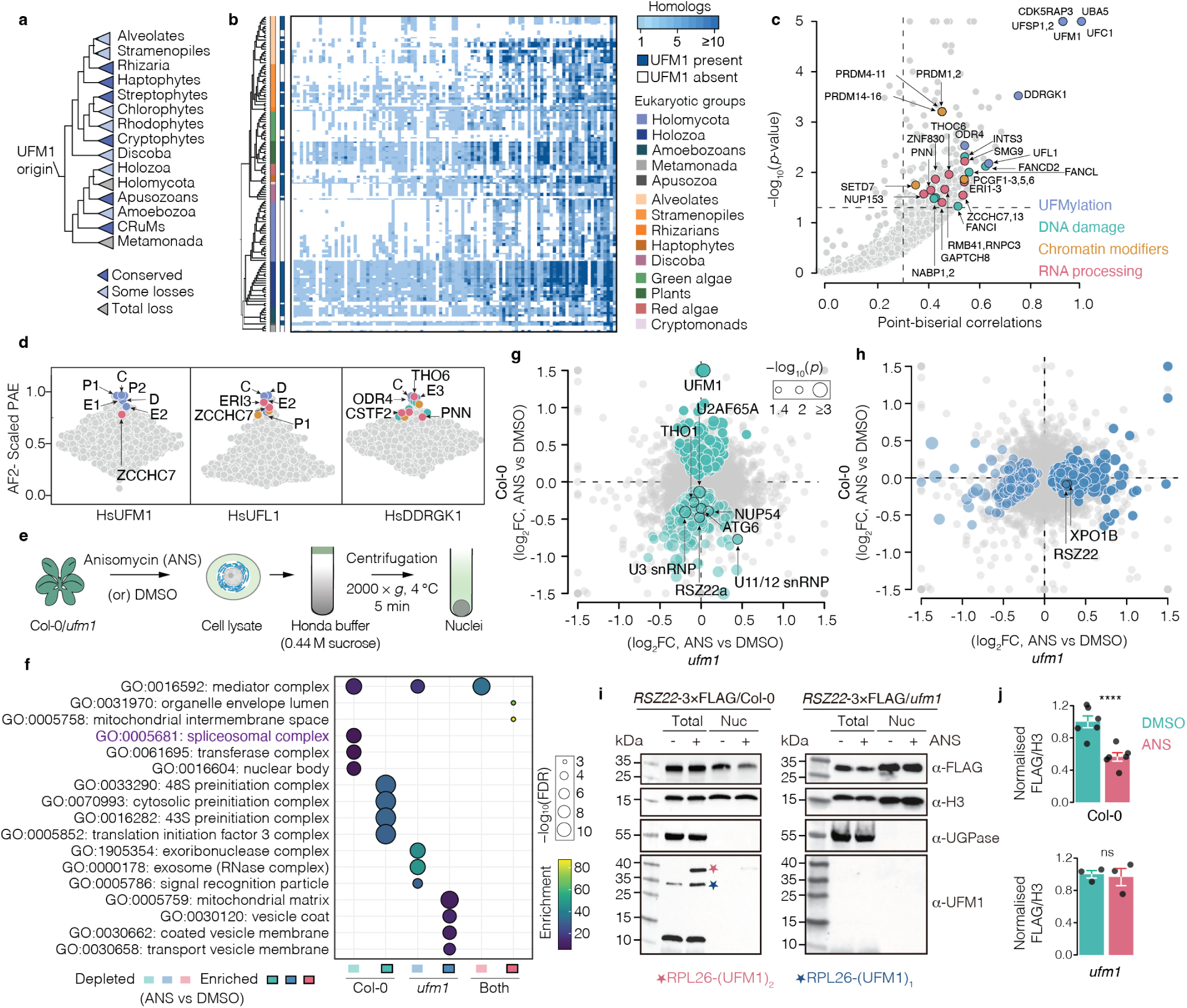
| Integrated evolutionary and proteomic profiling links UFMylation to nuclear mRNA processing. **a**, Phylogenetic distribution of UFM1 across major eukaryotic lineages. Conservation status is color-coded as conserved (blue), partial loss (light blue) or total loss (grey). The inferred evolutionary origin is indicated. **b**, Co-evolutionary heatmap of core UFMylation machinery and co-evolved genes across representative eukaryotes. Color intensity indicates copy number; supergroups shown on the right. **c**, Association plot of nuclear genes co-evolving with UFMylation components (point-biserial correlation *≥* 0.3, *P <* 0.05). Candidates are colored by functional category, highlighting enrichment in DNA damage, chromatin, and RNA processing pathways. **d**, Dot plot illustrating AlphaFold2 (AF2) Multimer interaction predictions of coevolved genes against UFMylation machinery. The Y-axis represents the average protein interaction score across the top three ranked models, calculated from domain-specific Predicted Aligned Error (PAE) values rescaled from 0 to 1. Abbreviations: P1, P2: UFSP1 and 2; E1, E2, E3: UBA5, UFC1, and UFL1; C: CDK5RAP3; D: DDRGK1. Coevolved genes with Scaled PAE *≥* 0.75 are colored according to key in Fig. 1c. **e**, Experimental workflow for nuclear fractionation of 10-day-old *Arabidopsis* Col-0 and *ufm1* seedlings treated with DMSO or anisomycin (ANS, 4 h). **f**, GO enrichment of proteins altered by ANS. Top four terms per condition (*≤* 300 genes, adjusted *P ≤* 0.01, fold enrichment *≥* 5) are shown. Dot size and color indicate *−* log_10_(FDR) and fold enrichment, respectively. **g, h**, Scatter plots of log_2_ fold-changes (ANS versus DMSO) in *ufm1* (x-axis) versus Col-0 (y-axis). Proteins significantly altered in Col-0 (green, **g**) or *ufm1* (blue, **h**) are highlighted, dot size reflects *−* log_10_(FDR). **i**, Immunoblotting analysis of RSZ22 abundance in total (whole cell lysates) and nuclear fractions from 10-day-old *RSZ22*-3*×*FLAG seedlings (Col-0 and *ufm1* backgrounds) treated with DMSO or ANS (4 h). H3 and UGPase are nuclear and cytosolic markers. Mono-(1) and di-(2) UFMylated RPL26 are indicated. See (Fig. S1m) and source data for replicates. **j**, Quantification of RSZ22 nuclear abundance normalized to H3. Dots represent independent biological replicates; bars indicate mean *±* standard error of the mean (s.e.m.). Statistical significance: Wilcoxon rank sum test, **** *P <* 0.0001; *ns*, not significant. **Fig. S1 on page 27**

To assess whether this evolutionary link reflects physical interactions, we used AlphaFold2-Multimer to screen for potential contacts between UFMylation machinery and co-evolving proteins. High-confidence models predicted an interaction between the E3 ligase component DDRGK1 and the transcription-export (THO/TREX) subunit THO6 (Fig. 1d, Table S2). Coimmunoprecipitation in *Arabidopsis* further confirmed that THO6 physically associates with both DDRGK1 and C53 (Fig. S1c, d).

Since our phylogenetic profiling and AlphaFold multimer analyses indicated a link between UFMylation and the nucleus, we next tested whether UFMylation remodels the nuclear proteome during translational stress. We performed quantitative proteomics on nuclear fractions from wild-type (Col-0) and *ufm1* mutant *Arabidopsis* seedlings subjected to translational stress (anisomycin, ANS) (Fig. 1e, Table S3). Fractionation purity was validated by Histone H3 enrichment and the absence of the cytosolic marker UGPase (Fig. S1e, f). Gene Ontology analysis revealed that spliceosome-related proteins were significantly reduced in the nuclear fraction in a UFM1-dependent manner (Fig. 1f). Consistent with phylogenetic predictions, proteomic profiling confirmed the reduced nuclear abundance of multiple mRNA processing factors, including spliceosomal components and splicing factors (e.g., RSZ22, RSZ22a), THO/TREX subunits (THO1), and nucleoporins (Fig. 1g, h). The known UFMylation-associated autophagy protein ATG6 was also depleted from the nuclear fraction^23^. Importantly, integrated proteomic and RNA-seq analysis indicated that these changes were independent of transcriptional regulation (Fig. S1g–l; Table S4). Since UFMylation is induced at the ER, we reasoned that nucleocytoplasmic shuttling proteins among the nuclear-depleted factors could represent candidates for UFM1-dependent spatial regulation. We focused on RSZ22 for further investigation, as it is a well-characterized member of the serine/arginine-rich (SR) protein family with established nucleocytoplasmic shuttling activity^33^. Immunoblotting validated the UFM1-dependent reduction of nuclear RSZ22 following stress, while total cellular levels in whole-cell lysates remained unchanged (Fig. 1i–j; Fig. S1m). Together, these data indicate that UFMylation regulates the stress-induced redistribution of mRNA processing factors between the nucleus and cytoplasm.

### UFMylation retains nucleocytoplasmic shuttling factors at the ER upon ribosome stalling

Given that SR proteins are evolutionarily conserved nucleocytoplasmic shuttling factors^34,35^, we hypothesized that UFMylated ribosomes at the ER could serve as platforms for spatially retaining these factors, thereby depleting them from the nucleus under stress. To test this, we isolated ER-derived microsomal fractions from wild-type (Col-0) and *ufm1* mutant *Arabidopsis* seedlings treated with DMSO or ANS (Fig. 2a). Microsomal purity was confirmed by enrichment of the ER marker CNX1/2 and the absence of the cytosolic protein NIT1 (Fig. S2a). Principal component analysis revealed that the microsomal proteome displayed clear segregation by genotype and stress treatment, whereas the nuclear proteome showed limited separation (Fig. 2b and Fig. S2b). This disparity likely reflects the fact that sequestration of shuttling factors at the ER, a compartment where they are normally sparse, generates a high-contrast signal, whereas their depletion from the nucleus represents a subtle shift against a large nuclear background.

**Figure 2.**
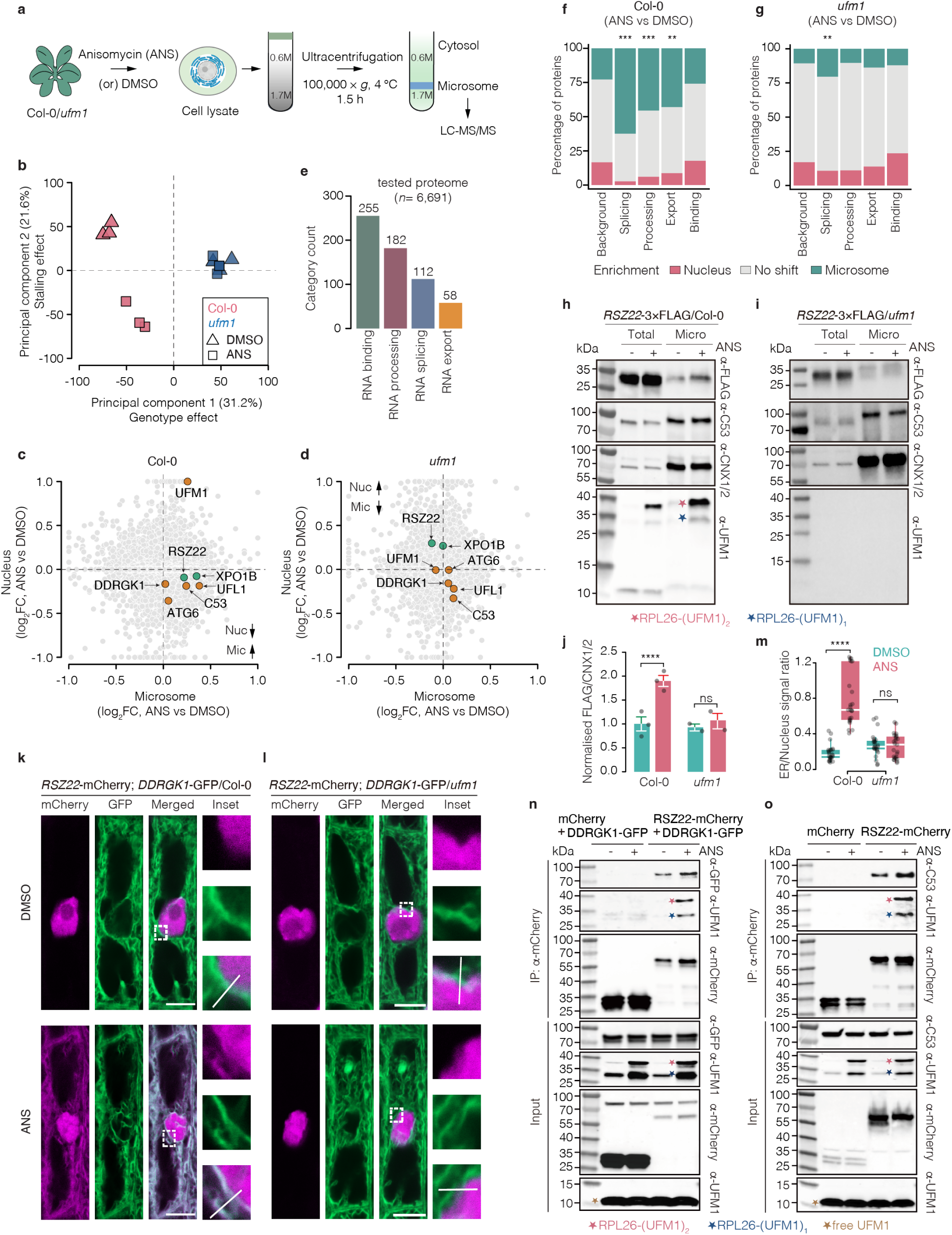
| UFMylation retains nucleocytoplasmic shuttling factors at the ER upon ribosome stalling. **a**, Experimental workflow for microsomal proteome analysis of 10-day-old seedlings treated with DMSO or ANS (4 h). **b**, Principal component analysis (PCA) of microsomal proteome (*n* = 3 biological replicates). PC1 and PC2 reflect proteomic separation by genotype and treatment. **c, d**, Scatter plots comparing ANS-induced log_2_ fold-changes in microsomal (x-axis) versus nuclear (y-axis) fractions in Col-0 (**c**) and *ufm1* (**d**). RNA-associated factors (e.g., RSZ22, XPO1B) and UFMylation machinery components are highlighted, illustrating UFM1-dependent redistribution. **e**, Functional annotation of quantified microsomal proteome (total quantified proteins *n* = 6, 691), showing number of RNA-related proteins by categories. **f, g**, Stacked bar charts show the distribution of proteins across subcellular enrichment classes following ANS treatment. Proteins were classified based on Δ enrichment, defined as log_2_FC(microsome) *−* log_2_FC(nucleus): Δ *>* +0.25 indi-cates microsome enrichment, Δ *< −*0.25 indicates nuclear enrichment, and intermediate values indicate no detectable shift. Statistical significance denotes enrichment of microsome-shifted proteins relative to all quantified proteins (back-ground), assessed using Fisher’s exact test with Benjamini-Hochberg FDR correction ( ** *P <* 0.01, *** *P <* 0.001). **h, i**, Immunoblot analysis of *RSZ22*-3*×*FLAG and endogenous C53 in total lysates and microsomal fractions (*n* = 3 for *RSZ22*-3*×*FLAG in Col-0; *n* = 2 for *RSZ22*-3*×*FLAG in *ufm1*). CNX1/2 serves as an ER marker. Mono-(1) and di-(2) UFMylated RPL26 are indicated. See (Fig. S2f–h) and source data for replicates. **j**, Quantification of RSZ22 microsomal enrichment (FLAG/CNX1/2 ratio) from (**h**), (**i**); Bars indicate mean *±* s.e.m. Statistical significance: Wilcoxon rank sum test, **** *P <* 0.0001; *ns*, not significant. **k, l**, Representative confocal images of RSZ22-mCherry and DDRGK1-GFP (ER marker) in root cells from Col-0 (**k**) and *ufm1* (**l**) backgrounds. ANS (1–2 h) induces RSZ22-ER signal overlap in Col-0 (**k**), and the signal overlap was absent in *ufm1* (**l**). Enlarged views and white lines indicate line-scan regions. Scale bars, 10 *µ*m. See (Fig. S3a–b) for line-scan profiles. **m**, Quantification of ER-to-nucleus RSZ22 signal ratios from (**k**), (**l**). Data are mean *±* SD from 3 biological replicates. Each dot represents an individual cell (*n* = 15 cells from 3 biological repli-cates). Bars indicate mean *±* SD. Statistical significance was determined by two-tailed unpaired t-test **** *P ≤* 0.0001; *ns*, not significant. **n, o**, In vivo Co-IP of RSZ22 interactions with DDRGK1 and UFMylated RPL26 (**n**) or interaction of RSZ22 with endogenous C53 and UFMylated RPL26 (**o**) under DMSO or ANS (4 h) treatment (*n* = 2). Proteins were extracted from *Arabidopsis* lines co-expressing DDRGK1-GFP with either mCherry (control) or RSZ22-mCherry in Col-0 background seedlings (**n**), or from mCherry (control) and RSZ22-mCherry seedlings (**o**). Input samples were normalized by Bradford assay prior to loading. The consistent level of endogenous free UFM1 (unconjugated) in input fractions serves as an internal loading control to show equal protein loading across all lanes. RSZ22-mCherry was immunoprecipitated using RFP-Trap beads. Mono-(1), di-(2) UFMylated RPL26 and free UFM1 are indicated. See (Fig. S3c–d) for replicates. **Fig. S2 and S3 on page 29 and 30**

Quantitative analysis of the microsomal proteome revealed significant UFM1-dependent enrichment of several nucleocytoplasmic shuttling factors following stress, including RSZ22 and the export factor XPO1B (Fig. 2c, d). RSZ22 is the *Arabidopsis* homolog of human SRSF7 and undergoes active shuttling between the nucleus and cytoplasm, a process partially mediated by the XPO1 export pathway^33,36,37^. Integrating the nuclear and microsomal datasets revealed a reciprocal redistribution pattern: multiple mRNA processing factors exhibited decreased nuclear abundance coupled with increased ER retention, and this redistribution was UFM1-dependent (Fig. 2e–g and Fig. S2d, e, Table S5). Immunoblotting confirmed this spatial shift for RSZ22, showing increased ER association with a corresponding reduction in nuclear levels following ANS treatment (Fig. 1i, j; Fig. 2h–j; Fig. S1m; S2f–h). Confocal imaging of RSZ22-mCherry with the ER marker DDRGK1-GFP confirmed a shift from nucleus to cytoplasm, partially localizing to the ER upon stress, which was absent in *ufm1* mutants (Fig. 2k–m and Fig. S3a, b).

We next tested whether this mechanism extends to other co-evolved factors. THO6, which physically associates with the UFMylation machinery (Fig. 1c and Fig. S1c, d), similarly accumulated at the ER in a UFM1-dependent manner upon stress (Fig. S3e–i). To determine whether the microsomal retention of RSZ22 is mediated by physical association with the UFMylation machinery, we performed co-immunoprecipitation experiments. RSZ22 associated with UFMylation components C53 and DDRGK1, as well as with UFMylated RPL26 (Fig. 2n, o and Fig. S3c, d), consistent with its ER recruitment. Together, these findings demonstrate that ribosome stalling induces UFM1-dependent retention of nucleocytoplasmic shuttling factors at the ER, providing a mechanism for their nuclear depletion.

### UFM1-dependent anchoring of splicing factors to stalled ER-associated ribosomes

To determine whether ER retention occurs directly at ribosomes, we performed proteomics of ribosomes isolated from *Arabidopsis* seedlings treated with ANS at various times (4 h, pooled time-course, and 16 h). Ribosomal fractions (60S, monosomes, and disomes) were isolated via sucrose-gradient fractionation and analyzed by LC-MS/MS (Fig. 3a). As an internal benchmark, the UFMylation E3 ligase complex (C53, UFL1, and DDRGK1) was enriched on stalled ribosomes in a UFM1-dependent manner (Fig. 3b and Fig. S4a–f, Table S6).

**Figure 3.**
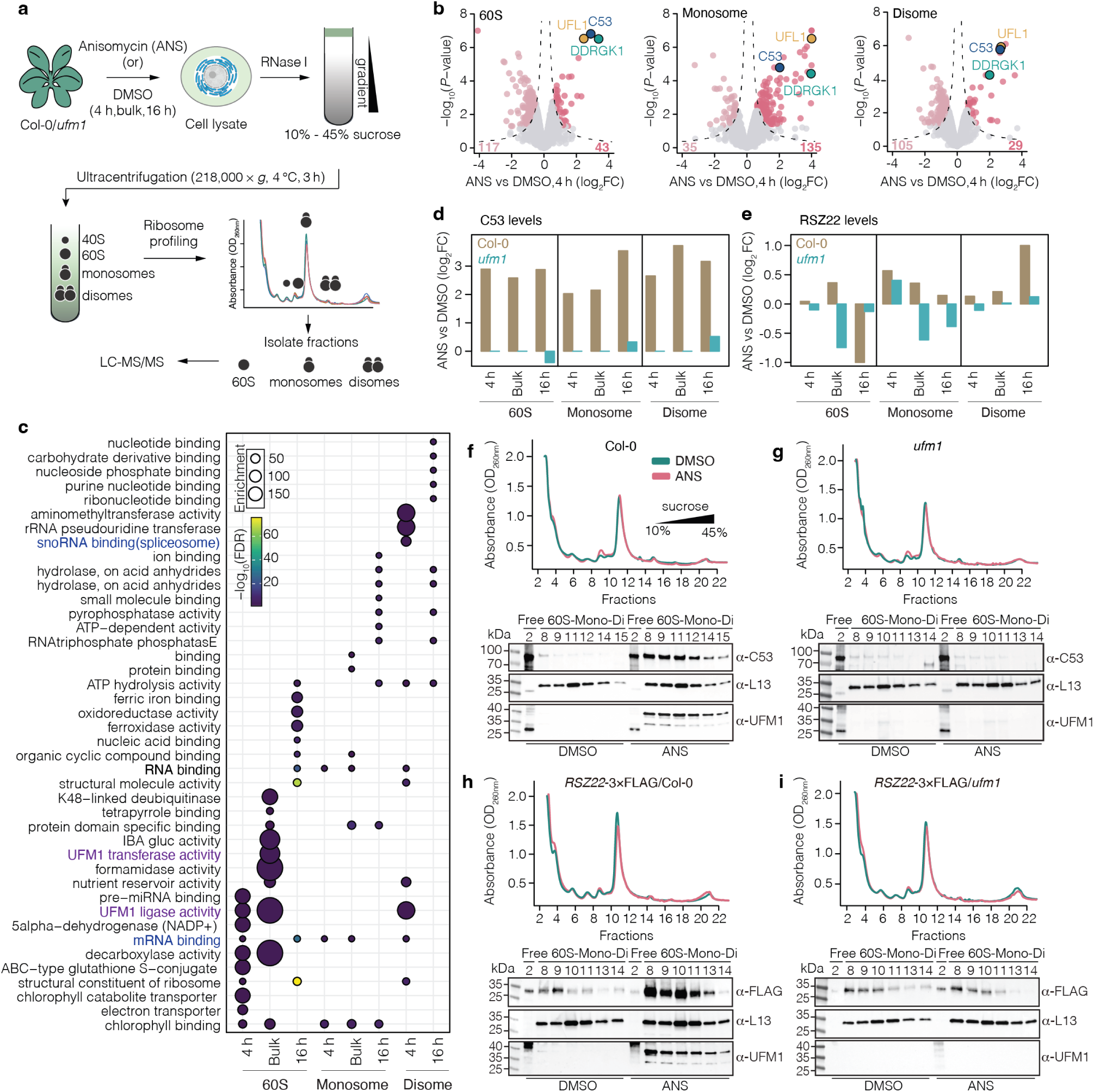
| UFM1-dependent anchoring of splicing factors to stalled ER-associated ribosomes. **a**, Schematic work-flow for ribosome proteomics using 10-day-old Col-0 and *ufm1* seedlings treated with DMSO or ANS. Ribosomal fractions (60S, monosome and disome) were isolated by RNase I digestion and sucrose gradient sedimentation for LC-MS/MS. Three datasets were analyzed: 4 h, a bulk sample (pooled from 1, 4, 8, 16 h treatments), and 16 h. **b**, Volcano plots (4-h dataset) showing ANS-induced recruitment of UFMylation components (UFL1, C53 and DDRGK1) to 60S, mono-somes and disomes in Col-0. x-axis, log_2_FC (ANS versus DMSO); y-axis, *−* log_10_(*P* value). **c**, GO enrichment analysis of stalling induced ribosome-enriched proteins in Col-0 plants. Dot color and size indicate *−* log_10_(FDR) and fold enrich-ment, respectively. Representative functional categories are highlighted. **d, e**, MS-based abundance profiles of C53 (**d**) and RSZ22 (**e**) across ribosomal fractions. Bars represent log_2_ fold change (ANS versus DMSO) in Col-0 (olive) and *ufm1* (cyan) from three datasets (4 h, bulk and 16 h). **f, g**, Sucrose gradient sedimentation and immunoblot analysis of endogenous C53 from Col-0 (**f**) and *ufm1* (**g**) seedlings treated with DMSO or ANS (4 h). UV absorbance (A_260_) profiles (top) indicate ribosome distribution across 10%–45% gradients. Fractions were immunoblotted for C53, L13 (60S marker), and UFM1 (*n* = 3). Free: ribosome-free fractions. See (Fig. S5d–g) for replicates. **h, i**, Sucrose gradient sedimentation and immunoblot analysis of *RSZ22*-3*×*FLAG in Col-0 (**h**) and *ufm1* (**i**) backgrounds. Fractions were immunoblotted for FLAG, L13, and UFM1 (*n* = 3). See (Fig. S5h–k) for additional replicates. **Fig. S4 and S5 on page 31 and 32**

Gene Ontology analysis of the stalling-induced ribosome-enriched proteins revealed the significant enrichment for mRNA binding and spliceosome proteins (Fig. 3c). Intersection analysis revealed a complex, translation-state-specific response to ANS stress (Fig. S4g). Heatmap clustering showed a largely intact global response in the *ufm1* mutant. But distinct subsets of candidate proteins showed UFM1-dependent association (Fig. S4h). For example, both C53 and the splicing factor RSZ22 are recruited to ribosomes in a stress and UFM1-dependent manner, while total cellular levels remained unchanged (Fig. 3d–i and Fig. S5a–c). Independent biological replicates confirmed the robustness of this association (Fig. S5d–k).

Importantly, the UFM1-dependent retention of mRNA processing factors represents a functionally distinct layer from known stalled ribosome sensing and splitting process. The conserved factors PEL1 and MBF1b (plant homologs of mammalian PELO and EDF1, respectively)^10,38,39^ were similarly enriched on stalled ribosomes in both wild-type and *ufm1* mutants (Fig. S4i, j). These results demonstrate that while the foundational ribosome rescue machinery remains intact in the *ufm1* mutant and operates independently of UFMylation, the UFM1 cascade supports an additional signaling function: capturing nucleocytoplasmic shuttling factors at ER-associated ribosomes. Together, these results establish that translational stress triggers UFM1-dependent recruitment of mRNA processing proteins to stalled ribosomes, where they are physically retained.

### UFM1-dependent intron retention selectively targets ER-related transcripts across species

To test the functional consequences of UFM1-dependent splicing factor retention, we analyzed stalling-induced alternative splicing across three systems: *Arabidopsis* seedlings, human colorectal carcinoma cell line (RKO), and mouse primary neurons. We fractionated the *Arabidopsis* and human samples into cytosolic and nuclear components, while total RNA was used for the mouse primary neurons. Ribosome stalling triggered widespread alternative splicing in all three models, with intron retention (IR) as the conserved outcome (Fig. 4a–c, Table S7). Strikingly, the magnitude of this response, measured by the mean absolute error (MAE) of aggregate splicing changes was significantly reduced in *ufm1* mutants, indicating that a substantial fraction of IR events depends on UFM1 (Fig. 4a–f and Fig. S6a–e, h–k). This establishes UFMylation as a key conserved regulator of stress-induced splicing via the spatial retention of splicing factors at ER-associated ribosomes.

**Figure 4.**
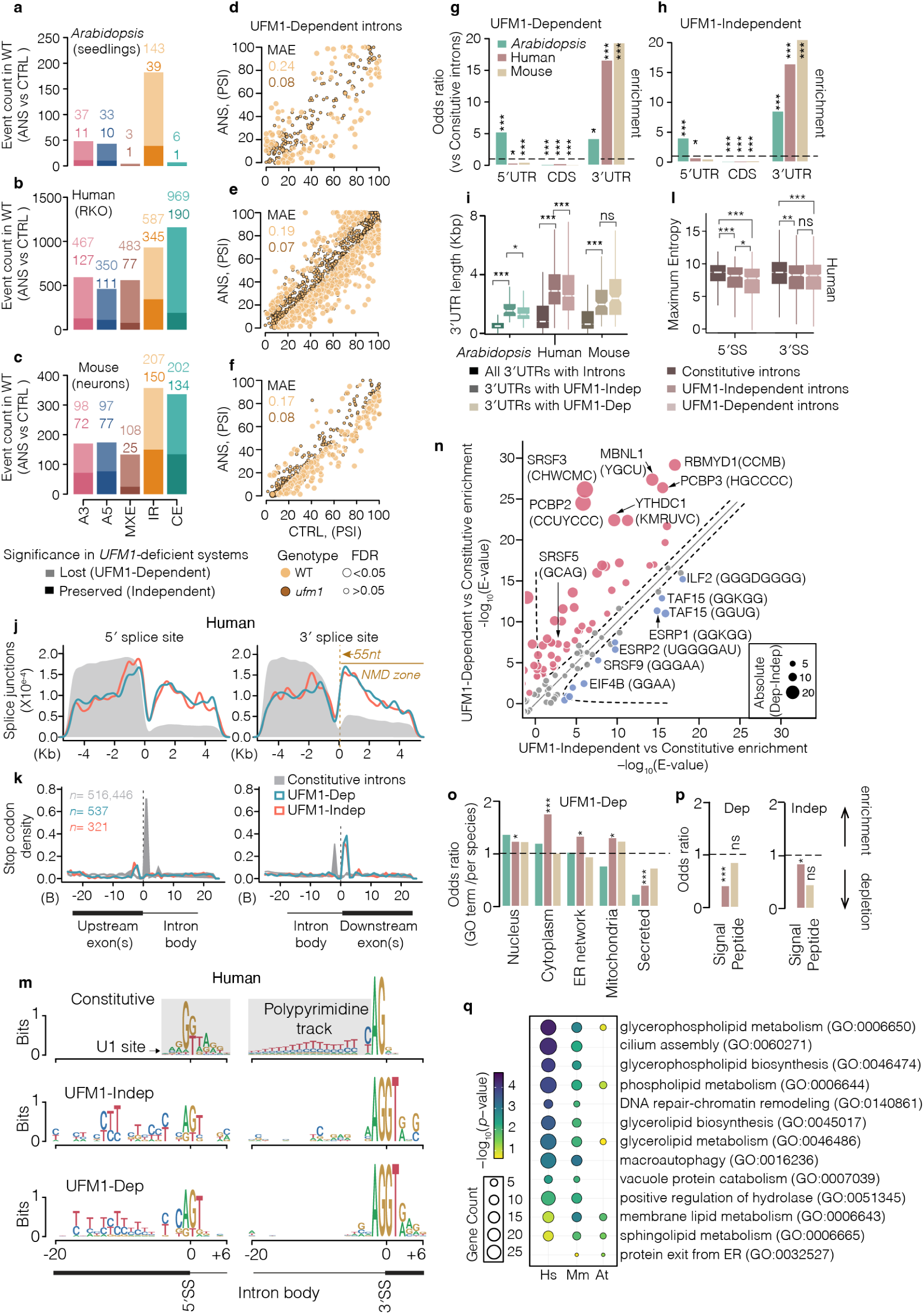
| A conserved UFM1-dependent program of intron retention targets a distinct RNA landscape. **a–c**, Quantification of alternative splicing changes induced by anisomycin (ANS)-mediated ribosome stalling in nuclear RNA from *Arabidopsis thaliana* seedlings (**a**) and human RKO cells (**b**), and total RNA from mouse neurons (**c**). Significant events were identified in wild-type (WT) samples using rMATS (FDR *<* 0.05) with an absolute change in percent spliced-in (*|*ΔPSI*| >* 0.1), for all event types except cassette exons (*|*ΔPSI*| >* 0.2). Events detected in WT but absent in *UFM1*-deficient systems (human *UFM1* knockout, mouse *Ufm1* knockout, and *Arabidopsis ufm1* knockout) were classified as UFM1-dependent (light shading); events in both WT and *UFM1*-deficient backgrounds were classified as UFM1-independent (dark shading). Event type: A3, alternative 3^′^ splice site; A5, alternative 5^′^ splice site; MXE, mutually exclusive exons; IR, retained intron; CE, cassette exon. **d–f**, Scatter plots comparing percent spliced-in (PSI) value under control and ANS conditions for UFM1 dependent intron retention events. WT events are shown in yellow, and events in *UFM1*-deficient systems are shown in dark yellow with black outlines; dot size reflects the False Discovery Rate (FDR). Mean Absolute Error (MAE) values summarize global splicing shifts per genotype. **g, h**, Genomic distribution of UFM1-dependent (**g**) and independent (**h**) retained introns across 5^′^ untranslated regions (UTRs), coding sequences (CDS), and 3^′^UTRs. Odds ratios calculated relative to constitutive introns. Ribosome stalling-dependent intron removal is significantly enriched in 3^′^ untranslated regions (3^′^UTRs) (Fisher’s exact test; \**P <* 0.05, \*\**P <* 0.001, \*\*\**P <* 0.0001). **i**, Length distribution of 3^′^UTRs colored by species and shaded by type: all 3^′^UTRs with introns (dark shade), 3^′^UTRs with UFM1-independent IRs (medium shade), and 3^′^UTRs with UFM1-dependent IRs (light shade). The 3^′^UTRs undergoing stalling-induced intron retention are significantly longer than all 3^′^UTRs with constitutive introns in the genome (Wilcoxon rank-sum test; \**P <* 0.05, \*\**P <* 0.001, \*\*\**P <* 0.0001). **j, k**, Metagene profiles of splice junction coverage (**j**) and stop codon density (**k**) relative to intron position. Compared with constitutive introns (grey), which are efficiently spliced and typically reside downstream of stop codons (canonical 3^′^UTR introns), UFM1-dependent (green) and independent (orange) introns retain junction coverage and are enriched for premature termination codons near the 3^′^ splice site. Persistence junction signal downstream of the 55-nucleotide nonsense-mediated decay (NMD) boundary (yellow line) indicates these introns are potentially sensitive to NMD. Kb: Kilobases; B: Bases. **l**, Splice site strength quantified by maximum entropy (MaxEnt) scores. Both UFM1-dependent and independent introns display weaker 5^′^ and 3^′^ splice sites compared with constitutive introns (Wilcoxon rank-sum test; \*\*\**P <* 0.0001), with UFM1-dependent introns exhibiting significantly weaker 5^′^ splice sites. **m**, Sequence logos of nucleotide composition surrounding the 5^′^ and 3^′^ splice sites. UFM1-dependent introns characterized by a conserved 5^′^ exonic CAG motif and C-rich sequences (see (Fig. S7e–g) for nucleotide probability). **n**, Comparison of RNA-binding protein (RBP) motif enrichment in UFM1-dependent versus UFM1-independent introns relative to constitutive introns from human. The diagonal indicates equal enrichment (*y* = *x*). Dashed curves denote hyperbolic significance thresholds (*y* = *x* + *c* + *k/x*, where *c* = 1.0 and *k* = 1.5). Dot size represents the absolute difference in *−* log_10_(E-value) between UFM1-dependent and UFM1-independent enrichments. UFM1-dependent introns are preferentially enriched for CAG and C-rich motif-binding RBPs (e.g., SRSF3, YTHDC1, SRSF5); UFM1-independent introns enriched for G-rich binding factors (e.g., ILF2, TAF15). **o, p**, Subcellular localization (**o**) and signal peptide (**p**) analyses of transcripts containing stalling-induced introns. Colored according to key in Fig. 4g. Both UFM1-dependent and independent targets are depleted for secretory pathway annotations and signal peptides (Fisher’s exact test; \**P <* 0.05, \*\*\**P <* 0.0001). **q**, GO enrichment analysis of UFM1-dependent transcripts shows overrepresentations of biological processes related to membrane lipid metabolism and endomembrane-associated processes rather than protein secretion. **Fig. S6 and S7 on page 33 and 35**

We next characterized the features of stalling-responsive introns. While approximately half of IR events were strictly UFM1-dependent (Fig. 4a–c, Table S7), the overall IR landscape exhibited conserved positional biases: retained introns were specifically located within 3^′^UTRs that are longer than the genomic average, a feature typical for nonsense-mediated decay (NMD) substrates^40,41^ (Fig. 4g–i). Specifically, these introns are followed by downstream splice junctions beyond the 55-nt boundary that triggers NMD, a pattern conserved across species (Fig. 4j, k and Fig. S7a–d). In both *Arabidopsis* and human cells, stress-induced IR events detected in the nuclear fraction were depleted from the cytosolic fraction, suggesting potential NMD targeting (Fig. S6f, g). Both UFM1-dependent and –independent introns exhibited weak splice site signatures, including lower entropy (maximum entropy score) at the 3^′^ splice site and reduced polypyrimidine tracts (Fig. 4l, m and Fig. S7e–i).

What distinguishes UFM1-dependent introns? Focusing on introns whose retention was reduced in *ufm1* mutants, we identified a distinct RNA signature: even weaker 3^′^ splice sites, a conserved 5^′^ exonic CAG motif, and CT-rich sequences (Fig. 4m and Fig. S7e–i). Motif analysis in human cells predicted that these features facilitate recognition by SR proteins including SRSF3 and SRSF5, as well as the m^6^A reader YTHDC1 (Fig. 4n), suggesting selective regulation of mRNA splicing in UFM1-dependent manner.

Functional annotation revealed that genes affected by UFM1-dependent IR events were enriched for membrane lipid metabolism and endomembrane-associated processes but depleted for secreted proteins (Fig. 4o–q). In contrast, UFM1-independent events were enriched for spliceosome and RNA-processing components (Fig. S7j, k). This functional divergence suggests that UFMylation imposes a selective layer of regulation on transcripts encoding ER-remodeling machinery.

Together, these results demonstrate that ribosome stalling induces a conserved, UFM1-dependent splicing program defined by a unique RNA signature and enriched for membrane-associated cellular processes. By sequestering splicing factors at the ER, UFMylation couples translational stress to nuclear mRNA processing, selectively reshaping the expression of membrane-associated genes, likely leading to membrane remodeling.

## Discussion

Our findings establish UFMylation as a retrograde signaling system that couples ER translational stress to nuclear gene expression. By anchoring SR splicing factors at stalled ribosomes, UFMylation creates a spatial regulatory axis between the ER and nucleus that reprograms the splicing landscape (Fig. 5). The conservation of this pathway from plants to mammals suggests it represents a fundamental adaptation for coordinating membrane homeostasis with translational surveillance. This mechanism operates in parallel to the canonical Unfolded Protein Response but acts post-transcriptionally rather than through transcriptional reprogramming. The conservation of this pathway from plants to mammals suggests it represents a fundamental adaptation for coordinating membrane homeostasis with translational surveillance. While the three UPR branches (IRE1, PERK, and ATF6) restore ER homeostasis primarily through transcriptional reprogramming or translational attenuation^42,43^, the UFMylation-SR axis enables post-transcriptional adjustment of gene expression. This distinction has important implications: whereas UPR activation requires sustained stress to trigger transcriptional programs, UFMylation-mediated splicing changes can occur rapidly upon ribosome stalling, potentially providing a faster adaptive response. The bidirectional intron retention we observe is consistent with altered nuclear availability of SR factors. SR proteins regulate alternative splicing in a concentration-dependent manner^34^, and UFM1-dependent ER retention of RSZ22 (Fig. 2–3) reduces its nuclear abundance, potentially shifting the stoichiometric balance among nuclear SR family members^44−46^. Consistent with this hypothesis, the enrichment of SR-binding motifs among UFM1-dependent targets supports a direct mechanistic link between SR protein sequestration and splicing outcomes (Fig. 4n).

**Figure 5.**
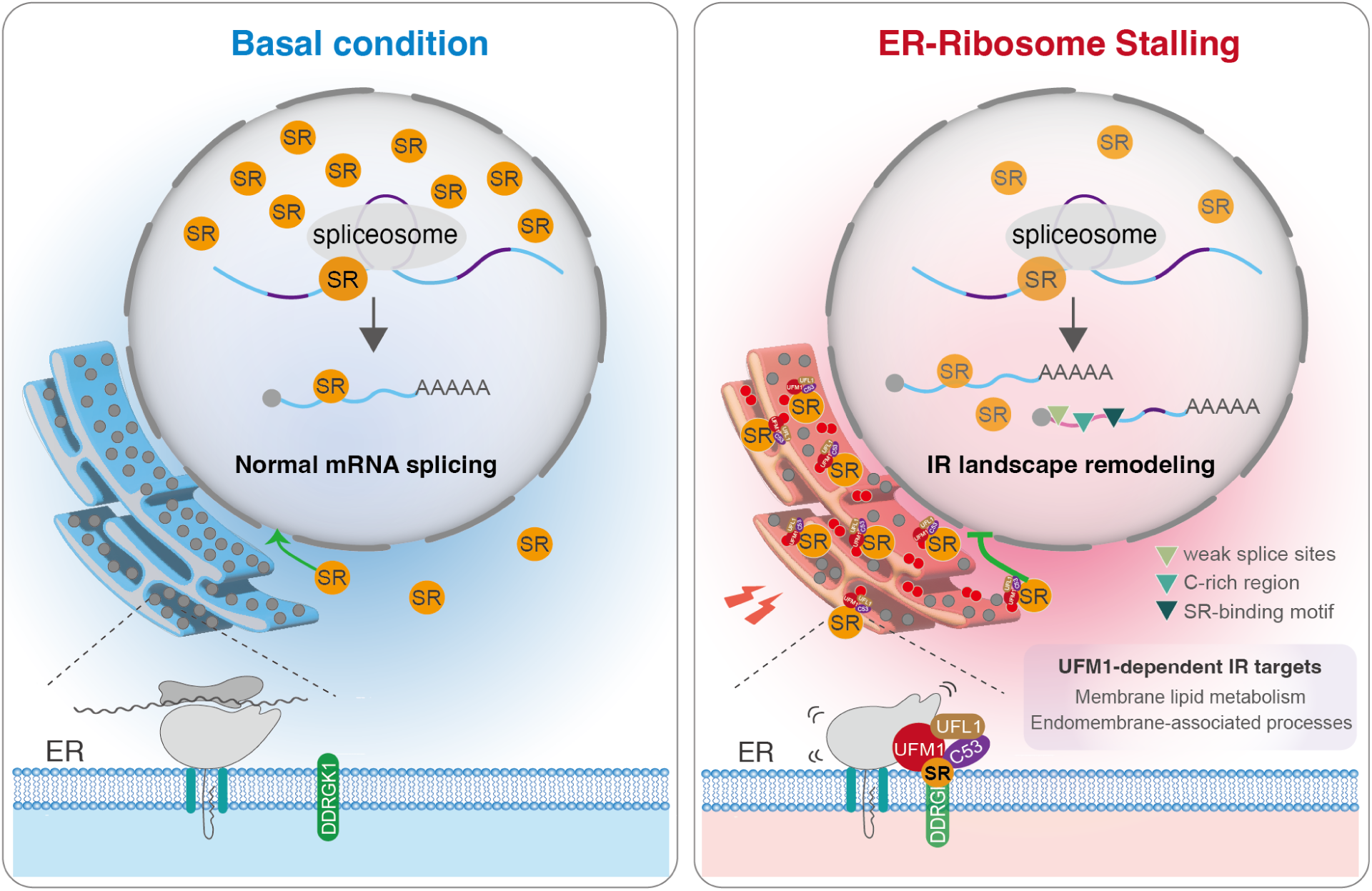
| Working model of UFM1-dependent spatial retention coupling ER-translational stress to nuclear splic-ing. Under basal conditions, SR shuttling proteins circulate between the ER and nucleus, supporting normal constitutive splicing. Under translational stress, activation of UFMylation at the translocon converts the ER-bound ribosomes into a docking platform, resulting in spatial retention of specific SR splicing factors (e.g., RSZ22) at the ER membrane, de-pleting their nuclear pool and remodeling the intron retention landscape of transcripts enriched for ER-related functions (e.g., membrane lipid metabolism and endomembrane-associated processes). UFM1-dependent introns are defined by a conserved RNA signature (weak splice sites, C-rich region, SR-binding motif). ER, endoplasmic reticulum; SR, serine/

The functional significance of UFM1-dependent splicing changes is underscored by convergent evidence across independent datasets. Cross-species analysis reveals that UFM1-responsive intron retention events are consistently enriched in pathways governing membrane lipid metabolism and endomembrane-associated processes (Fig. 4q). This functional bias is mirrored at the evolutionary level, where UFM1 co-evolves with a broad network of membrane-remodeling genes (Fig. S1a, Table S8), and at the proteomic level, where the stress-responsive microsome proteome is enriched for membrane remodeling factors (Fig. 2b, Fig. S2c, Table S9). These observations suggest that UFMylation does not primarily reduce protein load entering the ER, as might be expected from a quality control pathway, but rather reshapes ER membrane composition to adapt to translational stress. This reframing positions UFMylation as a coordinator of membrane homeostasis rather than simply a ribosome rescue mechanism.

Our findings also provide a mechanistic framework for understanding the unexpected connections between UFMylation and neurological diseases. The identification of UFMylation components as modifiers of tau aggregation in iPSC-derived neurons initially seemed paradoxical^29,30^, given tau’s cytosolic localization. Our discovery that UFMylation controls nuclear splicing through SR protein sequestration resolves this paradox: disruption of UFMylation would alter the splicing landscape of numerous transcripts, potentially including those encoding factors that influence tau proteostasis (Table S7). More broadly, the diverse pathologies associated with UFMylation defects^27,28,47,48^, including developmental abnormalities, metabolic dysfunction, and neurodegeneration, may reflect tissue-specific consequences of dysregulated splicing rather than solely defects in ribosome quality control. Our work reveals that ER-bound ribosomes serve not only as sites of protein synthesis and quality control but also as signaling platforms that communicate with the nucleus. The UFMylation-SR axis illustrates that organellar stress responses extend beyond local repair mechanisms to coordinate genome-wide changes in gene expression. While RSZ22 serves as a representative candidate, the extensive interactions within the SR family suggest it likely acts as part of a broader network subject to UFM1-dependent retention^33,49,50^. Whether other SR factors undergo similar retention and what molecular determinants govern their recognition remain key questions for future investigation. Given the essential role of UFMylation in development and its links to human disease, therapeutic modulation of this pathway, whether through targeting UFMylation itself or downstream splicing events, may offer new avenues for intervention in conditions characterized by ER stress.

## Materials and methods

### Plant materials and growth conditions

All *Arabidopsis thaliana* transgenic lines used in this study originate from the Columbia (Col-0) ecotype background and are listed in the materials section. Mutant lines used in this study are listed in the materials section.

Seeds were sterilized in 70% (v/v) ethanol with 0.5% (v/v) Tween-20 for 10 min, washed with absolute ethanol and sterile distilled water. After stratification at 4°C for 2 days in the dark, seeds were germinated in liquid half-strength Murashige and Skoog (1/2 MS) medium containing Murashige and Skoog salt + Gamborg B5 vitamin mixture, 1% (w/v) sucrose, and 0.5 g L^−1^ MES (pH 5.7). Seedlings were grown under a 16-h light/8-h dark photoperiod with a light intensity of 50 *µ*mol m^−2^ s^−1^ with constant shaking at 90 rpm.

For chemical treatments, 7– to 10-day-old *Arabidopsis* seedlings grown in liquid 1/2 MS medium were treated with 100 *µ*M anisomycin (ANS, Santa Cruz Biotechnology) or an equal volume of DMSO (control) for 4 h under continuous light with shaking at 90 rpm.

### Cloning and transgene construction

Coding sequences of *RSZ22* (At4g31580) and *THO6* (At2g19430) were amplified from Col-0 cDNA. PCR fragments were digested with *Bsa*I restriction enzyme and cloned into pMiniT 2.0 vectors with the same *Bsa*I site to generate intermediate constructs pMiniT-*RSZ22* and pMiniT-*THO6*. *Arabidopsis thaliana* transformation constructs were assembled using the GreenGate (GG) cloning system in the pGGSun backbone^51^. *E. coli* DH5*α* was used for plasmid propagation (Vienna BioCenter, in-house). All plasmids and primers used in this study are listed in the Materials section. Constructs were introduced into *Agrobacterium tumefaciens* strain GV3101 (C58 [RifR] Ti pMP90 [pTiC58DT-DNA] [gentR] Nopaline [pSoup-tetR]), all transgenic plants were generated using the floral dipping method^52^.

The *ufm1* mutant and the transgenic lines include pUbi::DDRGK1-GFP/Col-0 and pUbi::mCherry were described previously^4,53^. Double transgenic lines were generated by introducing constructs into existing transgenic backgrounds. Specifically, the pGGSun Ubi10::*RSZ22*-mCherry and pGGSun Ubi10::*THO6*-mCherry constructs were separately transformed into pUbi::DDRGK1-GFP/Col-0 plants to generate the pUbi::*RSZ22*-mCherry; pUbi::DDRGK1-GFP/Col-0 and pUbi::*THO6*-mCherry; pUbi::DDRGK1-GFP/Col-0 lines, respectively.

For experiments in the *ufm1* background, the pGGZ003::Ubi10 DDRGK1-GFP construct was introduced into pUbi::*RSZ22*-mCherry/*ufm1* and pUbi::*THO6*-mCherry/*ufm1* plants to obtain the respective double transgenic lines. Additionally, pUbi::DDRGK1-GFP; pUbi::mCherry double transgenic lines were generated by transforming the pGGZ003::Ubi10 DDRGK-GFP construct into pUbi::mCherry plants. All transgenic lines were confirmed by confocal microscopy or immunoblotting.

### Immunoblotting analysis in *Arabidopsis*

Total protein was extracted from 7-day-old *Arabidopsis* seedlings using Grinding Buffer (50 mM Tris-HCl, 150 mM NaCl, 1% Glycerol, 0.5% NP-40, 1.5 mM MgCl_2_, 1 × Protease inhibitor cocktail (Roche)). Lysates were incubated on ice for 10 min and cleared by two rounds of centrifugation (16,000 × *g*, 10 min, 4 °C). Protein concentration was determined via the Bradford method using Coomassie Blue G-250 (Sigma-Aldrich). Equal amounts of proteins were mixed with SDS loading buffer (250 mM Tris-HCl pH 6.8, 10% SDS, 0.05% bromophenol blue, 30% glycerol, freshly supplied with 5% of *β*-mercaptoethanol), boiled at 95 °C for 10 min. 10 *µ*g of total protein was loaded to the 4–20% Mini-PROTEAN TGX Precast Protein Gels (BioRad). Proteins were transferred to nitrocellulose membranes using a semi-dry Turbo Transfer Blot System (BioRad). Membranes were blocked with 5% skimmed milk in TBS and 0.1% Tween 20 (TBS-T) for 1 h at room temperature or overnight 4 °C and were subsequently incubated with primary antibody for 1 h at room temperature, followed by incubation with secondary antibody conjugated to horseradish peroxidase (HRP) for 1 h at room temperature. After three 10-min washes with TBS-T, signals were developed using Pierce ECL Western Blotting Substrate or ECL SuperSignal West Femto (Thermo Fisher Scientific) and visualized with a ChemiDoc Touch Imaging System (BioRad) or iBright Imaging System (Invitrogen). Images were processed using Image Lab software (BioRad).

Protein band intensities were quantified with ImageJ^54^. Rectangular regions of interest of equal size were defined around the target bands and the corresponding marker protein lanes (as internal controls for normalization). Band intensities were measured as the area under the peak from the generated profile. Target protein levels were normalized to the internal controls. Relative protein abundance was calculated from at least three biological replicates. The relative signal intensity of the control group samples was set as 1.0. The relative signal intensity of all other samples in each replicate was converted against the average signal intensity of the control sample. Data are presented as mean ± s.e.m.

Immunoblotting analyses were performed using the antibodies listed below, anti-AtC53, anti-GFP (Roche), anti-HsUFM1 (Abcam), anti-L13-1 (Agrisera), anti-UGPase (Agrisera), anti-CNX1/2 (Agrisera), anti-Nitrilase 1 (PhytoAB), anti-Histone H3 (Agrisera), anti-FLAG M2 (Sigma-Aldrich), and HRP-conjugated secondary antibodies (BioRad).

### Confocal microscopy and image analysis in *Arabidopsis*

Five-day-old *Arabidopsis* seedlings were used for confocal imaging. Seedlings were treated in liquid 1/2 MS supplemented with either DMSO (control) or 100 *µ*M anisomycin (ANS) for 1 h prior to microscopy. *Arabidopsis* roots were imaged using a Zeiss LSM 800 upright confocal microscope equipped with an Apochromat 40× objective lens (2× digital zoom). GFP was excited at 488 nm and detected at 510 nm; mCherry was excited at 561 nm and detected at 610 nm. All imaging parameters, including laser power, pinhole diameter, and detector gain, were kept constant across samples to ensure quantitative comparability.

Confocal images were processed using the ZEN (blue edition) and ImageJ. To evaluate the spatial relationship between target proteins and ER, fluorescence intensity line profiles were generated using the Plot Profile plugin in ImageJ. For each treatment, a linear ROI (Region of Interest) was drawn across representative cellular structures. The resulting gray values for each channel were plotted against distance (pixels). Colocalization was defined by the spatial coincidence of fluorescence intensity peaks across two channels. To quantify the ER/Nucleus signal ratio, three regions of interest (ROIs) were randomly selected within the nucleus and the ER. The mean gray value was calculated for each ROI to determine the relative signal intensity. For each cell, each of the three ER values was divided by all the nucleus values, and the mean of these three ratios was calculated as the representative signal ratio for that ROI. These three ROI-level ratios were then averaged to obtain a single value per cell. Quantification was performed on at least five representative images per treatment group, and the data are presented as mean ± s.e.m. from at least three biological replicates.

### *In planta* co-immunoprecipitation

For *in planta* Co-IP assays, 10-day-old *Arabidopsis* seedlings grown in liquid 1/2 MS medium were treated for 4 h in 1/2 MS liquid medium supplemented with DMSO (control) or 100 *µ*M anisomycin (ANS). Approximately 1–2 g plant material was harvested, flash-frozen in liquid nitrogen, and homogenized into a fine powder. Proteins were extracted using Co-IP grinding buffer (50 mM Tris-HCl pH 7.5, 150 mM NaCl, 10 mM MgCl_2_, 10% glycerol and 0.1% Nonidet P-40, 1× Protease inhibitor cocktail) by vortexing. Plant lysates were cleared by multiple rounds of centrifugation (16,000 × *g*, 5 min, 4 °C). To minimize non-specific binding, lysates were precleared by incubation with 10 *µ*L of Protein A Agarose (Roche) for 10 min at 4 °C. The protein concentrations were measured by Bradford assay. For each independent experiment, equal amounts of total protein across all samples and treatments were loaded for the Input analysis. For the pull-down, equal amounts of total proteins (2–3 mg) was incubated with 25 *µ*L of RFP-Trap Magnetic Agarose beads (ChromoTek) for 2.5 h at 4 °C with gentle rotation within each experiment. Beads were washed six times with Co-IP buffer and bound proteins were eluted by boiling the beads in SDS loading buffer for 10 min at 95 °C. The resulting supernatants were analyzed by immunoblotting as described above with respective antibodies. Endogenous unmodified UFM1 (free form) was detected on the same membrane in the input fractions as an internal loading control, as its level remained stable across all treatments.

### Subcellular fractionation from Arabidopsis

#### Microsome isolation

Ten-day-old *Arabidopsis* seedlings grown in liquid 1/2 MS medium were treated for 4 h in 1/2 MS liquid medium supplemented with DMSO (control) or anisomycin (ANS) and subsequently harvested for microsome isolation using a modified protocol^55^. Briefly, 1.5–2 g seedlings were homogenized in liquid nitrogen and resuspended in 6 mL of microsome extraction buffer (MEB; 100 mM Tris-HCl pH 8.0, 5 mM EGTA, 15 mM MgCl_2_, 2 *µ*M DTT, 0.3 M sucrose (Sigma-Aldrich), 20 U mL^−1^ SUPERase·In RNase Inhibitor (Thermo Fisher Scientific), 50 *µ* mL^−1^ cycloheximide (Sigma-Aldrich), and 1× Protease inhibitor cocktail). Homogenates were filtered with two layers of Miracloth and cleared by sequential centrifugation at 1,000 × *g* for 5 min and 8,000 × *g* for 10 min at 4 °C. Supernatant was layered on a step gradient (3 mL 0.6 M / 3 mL 1.7 M sucrose in MEB) in polyclear tubes (Laborgeraete Beranek), and centrifuged at 100,000 × *g* for 1.5 h at 4 °C using a TH-641 rotor (Sorvall Discovery SE Ultracentrifuge). The microsome fraction at the 0.6–1.7 M sucrose interface was collected into new polyclear tubes, diluted with 5 mL MEB buffer, and pelleted at 100,000 × *g* for 30 min at 4 °C to remove residue sucrose.

For mass spectrometry analysis, microsome pellets were precipitated following a trichloroacetic acid (TCA) precipitation method. Briefly, microsome pellets were resuspended in ultrapure water to a final volume of 1 mL, precipitated by adding 20 *µ*L of 10% sodium deoxycholate (NaDOC, Sigma-Aldrich) and 100 *µ*L of 50% TCA. Samples were vortexed thoroughly and incubated at −20 °C for 1 h to overnight. Proteins were pelleted by centrifugation at 16,000 × *g* for 30 min at 4 °C, washed with 1 mL of cold acetone, and pellets were air dried at room temperature. Protein pellets were further processed into peptides using the iST 96× kit (PreOmics) according to the manufacturer’s instructions.

For immunoblotting, microsome pellets were first resuspended in 0.5 mL of dissolving buffer (50 mM Tris–HCl pH 7.5, 150 mM NaCl, 10 mM MgCl_2_, 10% glycerol and 1% Nonidet P-40 supplemented with 1× Protease inhibitor cocktail). Pellets were homogenized by pipetting and incubated with gentle rotation at 4 °C for 20–30 min to ensure complete protein recovery. Proteins were subsequently subjected to TCA-precipitation as described above. The resulting protein pellets were fully dissolved in the same dissolving buffer. Protein concentrations were determined using the Pierce BCA Protein Assay Kits (Thermo Fisher Scientific), and the sample pH was adjusted with Tris-HCl (pH 7.5) to ensure proper electrophoretic migration. Equal amounts of microsome protein were denatured and resolved by SDS-PAGE for subsequent immunoblotting.

#### Nucleus isolation

Subcellular fractionation of nuclei was performed with modifications based on a previously described protocol^56^. Ten-day-old *Arabidopsis* seedlings (∼ 2g) grown in liquid 1/2 MS medium were treated for 4 h in the 1/2 MS liquid medium supplemented with DMSO or anisomycin. Seedlings were harvested, ground into fine powder in liquid nitrogen, and immediately resuspended in 5 mL of Honda buffer (0.44 M Sucrose, 1.25% Ficoll (Carl ROTH), 2.5% Dextran T40 (PanReac AppliChem), 20 mM HEPES, pH 7.4, 10 mM MgCl_2_, 0.5% Triton X-100, 1 mM DTT, 1 × Protease inhibitor cocktail and RNaseOUT nuclease inhibitor (Thermo Fisher Scientific)). The homogenate was filtered through two layers of Miracloth and centrifuged at 2,000 × *g* for 5 min at 4 °C to pellet nuclei. Supernatant was further centrifuged at 14,000 × *g* for 5 min at 4 °C. This step was repeated twice to remove residual nuclear contaminants, and the final supernatant was collected as the cytoplasmic fraction.

For isolation of the nuclear fraction, the pellet obtained after the initial 2,000 × *g* centrifugation was resuspended in 15 mL Honda buffer and centrifuged again at 2,000 × *g* for 5 min at 4 °C. The pellet was resuspended and washed once more with 10 mL of Honda buffer and centrifuged. The final nuclear pellet was resuspended in 1 mL of Honda buffer and transferred to a new RNase/DNase-free 1.5 mL microcentrifuge tube, followed by centrifugation at 8,000 × *g* for 1 min at 4 °C. The resulting pellet was resuspended in 500 *µ*L of Honda buffer for subsequent mass spectrometry and immunoblotting analyses. For nucleus mass spectrometry analysis, samples were processed using the TCA precipitation method followed by peptide purification using the iST 96× kit (PreOmics), as described above for microsome isolation.

#### Sucrose gradient fractionation of ribosomes for biochemical analysis in Arabidopsis

Sucrose gradient fractionation was performed with modifications based on previously described methods^39,57^. Seven-day-old *Arabidopsis* seedlings (∼ 0.5 g) grown in liquid 1/2 MS medium were harvested and ground into a fine powder in liquid nitrogen. The powder was resuspended in 150 *µ*L of protein extraction buffer (PEB; 0.1 M Tris–HCl pH 8.0, 40 mM KCl, 2.5 mM MgCl_2_, 0.4% polyoxyethylene-10-tridecyl ether (PTE, Sigma-Aldrich), 0.4% detergent mix (20% Triton, 20% Tween-20, 20% Brij-35 (Sigma-Aldrich), 0.4% sodium deoxycholate, 1 mM DTT, 100 *µ*g mL^−1^ cycloheximide). Samples were incubated on ice for 10 min with constant vortex to ensure resuspension. Cell debris was removed by centrifugation at 21, 000 × *g* for 10 min at 4 °C. The supernatant was transferred to a new tube, and the clarification step was repeated three times. Equal amounts of RNA (∼ 200 *µ*g) were adjusted to a final volume of 220 *µ*L. RNase I (Ambion, Thermo Fisher Scientific) was added at a ratio of 1 *µ*L per 100 *µ*g of total RNA, and samples were digested for 1 h 15 min at 4 °C with gentle rotation. Digestion was terminated by centrifugation and addition of 0.5 *µ*L SUPERase•In RNase Inhibitor.

A total of 200 *µ*L of each sample was loaded onto 10–45% (w/v) sucrose gradient prepared in gradient buffer (24 mM Tris–HCl pH 8.0, 16 mM Tris–HCl pH 9.0, 40 mM KCl, 10 mM MgCl_2_, 1 mM DTT and 100 *µ*g mL^−1^ cycloheximide). Gradients were centrifuged at 35, 000 rpm for 3 h at 4 °C using an SW-40 Ti rotor (Beckman Coulter Optima L-90K ultracentrifuge) or at 35, 700 rpm for 3 h at 4 °C using a TH-641 rotor (Sorvall Discovery SE ultracentrifuge).

Gradients were fractionated using a top-down Biocomp Piston Gradient Fractionator (Biocomp) equipped with a Triax UV detector and flow cell. UV (A_260_) absorbance across 10–45% sucrose gradients was continuously monitored, and fractions (0.5 mL per tube) were collected and stored on ice or flash-frozen in liquid nitrogen until further processing. Peaks corresponding to the 60S subunit, monosome, and disome were collected for subsequent mass spectrometry or immunoblotting analyses.

For ribosome-associated mass spectrometry, individual or pooled fractions corresponding to 60S, monosome, or disome peaks were flash frozen in liquid nitrogen and processed using TCA precipitation, followed by tryptic digestion and purification using the iST 96× kit, as described above. For ribosome SDS-PAGE and immunoblotting, each 0.5 mL fraction was adjusted to a final volume of 1 mL with ultrapure water and subjected to TCA-precipitation. Protein pellets were resuspended and denatured. Equal volumes of ribosome-containing fractions were loaded for immunoblotting, whereas ribosome-free fractions (free) were loaded at one-fifth of the volume used for the ribosome-containing fractions.

### Mass spectrometry data acquisition and analysis

All samples submitted for mass spectrometry were prepared using the iST 96× kit. 16 h and time-course pooled ribosome mass spectrometry samples, and the nucleus mass spectrometry samples were processed using the Orbitrap Exploris 480 mass spectrometer. 4 h ribosome mass spectrometry samples were processed using the Eclipse mass spectrometer. Orbitrap Exploris 480 and Eclipse mass spectrometer was operated in data-dependent mode, performing a full scan. Microsome mass spectrometry samples were processed using the timsTOF HT mass spectrometer. The timsTOF HT was operated in DIA mode where 1 MS1 scan was followed by 8 DIA-PASEF ramps.

#### nanoLC-MS/MS Analysis for data-dependant acquisition (DDA)

The nano HPLC system (UltiMate 3000 RSLC nano system or Vanquish Neo UHPLC-System, Thermo Fisher Scientific) was coupled to an Orbitrap Exploris 480 mass spectrometer equipped with a FAIMS pro interfaces and a Nanospray Flex ion source (all parts Thermo Fisher Scientific). Peptides were loaded onto a trap column (PepMap Acclaim C18, 5 mm × 300 *µ*m ID, 5 *µ*m particles, 100 Å pore size, Thermo Fisher Scientific) at a flow rate of 25 *µ*l min^−1^ using 0.1% TFA as mobile phase. After 10 minutes, the trap column was switched in line with the analytical column (PepMap Acclaim C18, 500 mm × 75 *µ*m ID, 2 *µ*m, 100 Å, Thermo Fisher Scientific or *µ*Pac 110cm generation 2 micropillar column, Pharmafluidics) operated at 30 °C (PepMap) or 50 °C (*µ*Pac). Peptides were eluted using a flow rate of 230 nl/min, starting with the mobile phases 98% A (0.1% formic acid in water) and 2% B (80% acetonitrile, 0.1% formic acid) and linearly increasing to 35% B over the next 180 min.

The Orbitrap Exploris 480 mass spectrometer was operated in data-dependent mode, performing a full scan (*m/z* range 350–1200, resolution 60,000, normalized AGC target 100%) at 3 different compensation voltages (CV –45, –60, –75), followed each by MS/MS scans of the most abundant ions for a cycle time of 0.8 seconds per CV. MS/MS spectra were acquired using HCD collision energy of 30, isolation width of 1.0 or 1.2 *m/z*, orbitrap resolution of 15,000, normalized AGC target 100%, minimum intensity of 25,000 and maximum injection time of 30 ms. Precursor ions selected for fragmentation (include charge state 2–6) were excluded for 40 s. The monoisotopic precursor selection filter and exclude isotopes feature were enabled.

The Eclipse was operated in data-dependent mode, performing a full scan (*m/z* range 380–1500, Orbitrap resolution 120k, AGC target value 400,000, normalized AGC target 100%) at 4 different compensation voltages (CV –45, –55, –65, –75), followed each by MS/MS scans of the most abundant ions for a cycle time of 0.6 sec per CV. MS/MS spectra were acquired using an isolation width of 1.2 *m/z*, absolute AGC target value of 10,000, normalized AGC target 100%, minimum intensity of 5,000 and a maximum injection time of 35 ms, using the Iontrap for detection with HCD collision energy of 30. Precursor ions selected for fragmentation (including charge state 2–6) were excluded for 20 s. The monoisotopic precursor selection filter and exclude isotopes feature were enabled.

#### Data Processing protocol for DDA Data

For peptide identification, the RAW-files were loaded into Proteome Discoverer (version 2.5.0.400, Thermo Scientific). All MS/MS spectra were searched using MSAmanda v2.0.0.19924^58^. The peptide mass tolerance was set to ±10 ppm and fragment mass tolerance to ±10 ppm for measurements done on the Exploris and to ±400 mmu for measurements done on the Eclipse, the maximum number of missed cleavages was set to 2, using tryptic enzymatic specificity without proline restriction. Peptide and protein identification was performed in two steps. For an initial search the RAW-files were searched against the *Arabidopsis* taxonomy database (TAIR; 32,785 sequences; 14,482,855 residues), supplemented with common contaminants and sequences of tagged proteins of interest using Iodoacetamide derivative on cysteine as a fixed modification. The result was filtered to 1% FDR on protein level using the Percolator algorithm integrated in Proteome Discoverer^59^. A sub-database of proteins identified in this search was generated for further processing. For the second search, the RAW-files were searched against the created sub-database using the same settings as above and considering the following additional variable modifications: oxidation on methionine, deamidation on asparagine and glutamine, glutamine to pyro-glutamate conversion at peptide N-terminal glutamine and acetylation on protein N-terminus. For some files phosphorylation on serine, threonine and tyrosine was included as variable modification in the search. The localization of the post-translational modification sites within the peptides was performed with the tool ptmRS, based on the tool phosphoRS^60^. Identifications were filtered again to 1% FDR on protein and PSM level, additionally an Amanda score cut-off of at least 150 was applied. Proteins were filtered to be identified by a minimum of 2 PSMs in at least 1 sample. Protein areas have been computed in IMP-apQuant by summing up unique and razor peptides^61^. Resulting protein areas were normalized using intensity-based absolute quantification (iBAQ) and sum normalization was applied for normalization between samples^62^. Match-between-runs (MBR) was applied for peptides with high confident peak area that were identified by MS/MS spectra in at least one run.

Proteins were required to be identified by a minimum of 2 PSMs in at least 1 sample, and quantified proteins were filtered to include at least 3 quantified peptide groups. The statistical significance of differentially expressed proteins was determined using the limma package^63^.

#### LC-MS/MS setup for Data independent Acquisition (DIA)

Peptide samples were analyzed by the LC-MS/MS approach. The nano HPLC system used was an UltiMate 3000 nano HPLC RSLC (Thermo Scientific) equipped nano-electrospray source (CaptiveSpray source 1, Bruker Daltonics), coupled to a timsTOF HT mass spectrometer (Bruker Daltonics).

Peptides samples were injected on a pre-column (PepMap C18, 5 mm × 300 *µ*m × 5 *µ*m, 100 Å pore size, Thermo Scientific) with 2% ACN/water (v/v) containing 0.1% TFA at a flow rate of 10 *µ*l min^−1^ for 10 min. Peptides were then separated on the 25 cm Aurora ULTIMATE series HPLC column (CSI; 25 cm × 75 *µ*m ID, 1.7 *µ*m C18, IonOpticks) operating at 50 °C controlled by the Column Oven PRSO-V1-BR (Sonation), using UltiMate 3000 (Thermo Scientific Dionex). The analytical column flow was set to 300 nL/min and the mobile phases water/0.1% FA and 80% ACN/0.08% FA (A and B, respectively) were applied in the linear gradients starting from 2% B: 2%–10% B in 10 min, 10%–24% B in 35 min, 24%–35% B in 15 min, 35%–95% B in 1 min, 95% for 5 min, and finally the column was equilibrated in 2% B for next 10 min (all % values are v/v; water and ACN solvents were purchased from Thermo Scientific at LC-MS grade).

The LC system was coupled with TIMS quadrupole time-of-flight mass spectrometer (timsTOF HT, Bruker Daltonics) and samples were measured in dia-PASEF mode. The CaptiveSpray source parameters were: 1600 V of capillary voltage, 3.0 L min^−1^ of dry gas, and 180 °C of dry temperature. MS data was acquired in the MS scan mode, using positive polarity, 100–1700 *m/z* range, mobility range was set up from 0.64–1.42 V·s/cm^2^, ramp time was set to 100 ms and estimated cycle time was 0.95 s. Collision energy was 27 eV at 1*/k*_0_ 0.85 V·s/cm^2^, and 45 eV at 1*/k*_0_ 1.3 V·s/cm^2^. Automatic calibration of ion mobility was set ON. The timsTOF HT was operated in DIA mode where 1 MS1 scan was followed by 8 DIA-PASEF ramps.

#### Proteomics data analysis for DIA data

DIA data was analysed in Spectronaut 18.7^64^. Trypsin/P was specified as a proteolytic enzyme and up to 2 missed cleavages were allowed in the Pulsar directDIA+ search. Dynamic mass tolerance was applied for calibration and main search. The search was performed against the *Arabidopsis* taxonomy database (TAIR; 32,785 sequences; 14,482,855 residues), with common contaminants (347 sequences) and common tags (28 sequences) appended. Here, Carbamidomethylation of cysteine was searched as fixed modification, whereas oxidation of methionine and acetylation at protein N-termini were defined as variable modifications. Peptides with a length between 7 and 52 amino acids were considered and results were filtered using Spectronaut default filtering criteria (Precursor Qvalue *<* 0.01, Precursor PEP *<* 0.2, Protein Qvalue *<* 0.01 per Experiment and *<* 0.05 per Run, Protein PEP *<* 0.75). Quantification was performed as specified in Biognosys BGS Factory Default settings, grouping Peptides by Stripped Sequence and performing protein inference using IDPicker. Cross-Run Normalization in Spectronaut was deactivated due to subsequent mode normalization.

Spectronaut results were exported using Pivot Reports on the Protein and Peptide level and converted to Microsoft Excel files using our in-house software MS2Go. For DIA data MS2Go utilizes the python library msReport (developed at the Max Perutz Labs Proteomics Facility) for data processing. Abundances were normalized by the mode of protein ratios in msReport and missing values were imputed with values obtained from a log-normal distribution with a mean of 100. To compensate for different protein lengths, protein quantification was then normalized using iBAQ^62^. Statistical significance of differentially expressed proteins was determined using limma^63^.

### Subcellular enrichment shifts quantification and functional categorization from Arabidopsis mass spectrometry data

To quantify subcellular enrichment shifts in response to anisomycin treatment, we compared log_2_ fold changes in normalized protein abundance between microsomal and nuclear fractions in both wild-type and *ufm1* mutant backgrounds. Only proteins detected in both compartments were retained to ensure comparability. For each protein, enrichment was defined as the difference in anisomycin-induced fold change between compartments (Anisomycin vs DMSO), calculated as: Δ enrichment = Microsome – Nuclear. This was computed separately for wild-type and *ufm1* mutants. To limit the influence of outliers, Δ enrichment values were winsorized to the range of −1 to +1. Proteins were categorized as microsome-enriched (Δ *>* 25%), nucleus-enriched (Δ *<* −25%), or no-shift (−25% ≤ Δ ≤ 25%).

Gene Ontology (GO) annotations were retrieved from Ensembl Plants using the biomaRt package and filtered for RNA-related terms containing keywords including splicing, binding, export, decay, and processing. These GO terms were grouped into six RNA-related categories: RNA binding, RNA splicing, RNA processing, RNA decay and stability, RNA export, and RNA catabolic processes. Functional categories with fewer than ten proteins were not considered for statistical testing. For each category, the proportion of proteins in each enrichment class was calculated and visualized using stacked bar plots. Fisher’s exact tests were used to assess whether category-specific enrichment patterns significantly deviated from the total proteome, and *P* –values were adjusted for multiple testing using the Benjamini-Hochberg method. In parallel, empirical cumulative distribution functions (ECDFs) of Δ enrichment values were generated using the base R ecdf() function to compare the distributional behavior of each RNA-related category against the global proteome. This allowed systematic evaluation of how protein localization patterns across RNA-associated functions are altered by stress and UFM1 pathway disruption.

### Human cell culture and cell line generation

RKO-iCas9 cells (doxycycline-inducible Cas9) were cultured in RPMI-1640 media (Sigma) supplemented with 10% fetal bovine serum (Gibco), 2 mM L-Glutamine (Sigma), 1% Penicillin-Streptomycin (Sigma), 1× Non-Essential Amino Acids (Thermo Scientific), and 1 mM sodium pyruvate (Gibco). Cell lines were grown in a humidified incubator at 37 °C and 5% CO_2_ and regularly tested for *Mycoplasma* infection with the EZ-PCR™ Mycoplasma Detection Kit (Biological Industries).

RKO-iCas9 cells expressing sgRNAs against the non-coding locus *AAVS1* were generated in^65^ and were a kind gift from the Gijs Versteeg lab. RKO-iCas9 cells expressing sgRNAs against *UFM1* were generated in our previous study^19^; gRNA sequences 5^′^-CGACGTCAGCGTGATCTTAA-3^′^ and 5^′^-ACACTTTGTACGGCAGCCGT-3^′^ were used to target the *UFM1* gene. The parental RKO-iCas9-GFP cell line was a kind gift from the Johannes Zuber lab^66^.

### Cytoplasmic and nuclear fractionation of human cells for RNA and protein analysis

0.75 million RKO-iCas9 cells were plated per 6-well plate. To induce Cas9 expression, 250 ng mL^−1^ doxycycline was added at the time of seeding. DMSO in 1:2000 dilution was used as a control. 72 h after doxycycline induction, cells were treated with 200 nM anisomycin (ANS) for 1 h followed by fractionation of cytoplasmic and nuclear RNA according to the published protocol^67^ with modifications. Cells were washed with ice-cold phosphate-buffered saline (PBS) and lysed on the plate in 335 *µ*L of 0.1% NP-40 in 1× PBS. Plates were incubated on ice for 5 min and lysates collected by scraping. Samples were pipetted 5 times, transferred to 1.5 mL Eppendorf tubes, and pipetted 2 additional times. 30 *µ*L were transferred to a separate tube for immunoblotting analysis (whole cell lysate).

Cell lysates were centrifuged at 3, 000×*g* for 2 min at 4 °C. 250 *µ*L of supernatant were transferred to 1.5 mL Eppendorf tubes for a second round of centrifugation at 3, 000 × *g* for 2 min at 4 °C. 130 *µ*L of supernatant (cytosolic fraction) were transferred to 1.5 mL Eppendorf tubes. 30 *µ*L were transferred to a separate tube for immunoblotting analysis and 100 *µ*L were kept for RNA extraction. The nuclear pellet was washed with 1 mL of 0.1% NP-40 in 1× PBS and centrifuged at 3, 000 × *g* for 2 min at 4 °C. After removal of the supernatant, the pellet was used for RNA extraction or dissolved in 1× SDS sample buffer (50 mM Tris-HCl pH 6.8, 2% SDS, 0.1% Bromophenol blue, 10% glycerol, 20 mM DTT) for immunoblotting analysis. The following antibodies were used for immunoblotting: anti-UFM1 (Abcam) at 1:2000, anti-lamin A/C (Santa Cruz) at 1:1000, anti-eEF2 (Proteintech) at 1:10000, and IgG (H+L), HRP Conjugates (Promega).

### Mouse lines and primary hippocampal neuronal culture

Mouse lines were maintained in the animal facility of the Max Planck Institute for Multidisciplinary Sciences. The *Ufm1*-cKO mouse line, which carries loxP sites 5^′^ of exon 3 and 3^′^ of exon 5 of the *Ufm1* gene, was obtained from GemPharmatech. The physiological consequences of Cre-induced *Ufm1*-loss in cultured neurons were described previously^28^. The line was kept in a homozygous mutant state. Animals homozygous for *Ufm1*-cKO were viable, fertile, and exhibited no overt phenotypic changes in the cage environment.

We have complied with all relevant ethical regulations for animal use and received ethical approval from the Niedersächsisches Landesamt für Verbraucherschutz und Lebensmittelsicherheit (LAVES). Mouse breeding was done with permission of the LAVES and according to European Union Directive 2010/63/EU. Mice were housed in individually ventilated cages with bedding and nesting material in specific pathogen-free conditions at 21 °C and 55% relative humidity, under a 12 h/12 h light/dark cycle. Mice received food and water *ad libitum*. Health monitoring was done quarterly following FELASA recommendations with either NMRI sentinel mice or animals from the colony. All mice used for experiments were sacrificed by decapitation on their day of birth. Animals from both sexes were used without further distinctions.

Mouse primary hippocampal neuron cultures were prepared as described previously^28,68,69^. Briefly, brains of newborn mice were dissected, and hippocampi were isolated and digested for 1 h at 37 °C in Dulbecco’s modified Eagle’s medium (DMEM) containing 2.5 U mL^−1^ papain (Worthington Biomedical Corporation), 0.2 mg mL^−1^ L-cysteine, 1 mM CaCl_2_, and 0.5 mM EDTA. Papain digestion was stopped by incubation with DMEM supplemented with 10% (v/v) heat-inactivated fetal bovine serum (FBS), 2.5 mg mL^−1^ albumin, and 2.5 mg mL^−1^ trypsin inhibitor. Hippocampi were triturated in Neurobasal medium (Neurobasal medium-A) containing 2% (v/v) B27, 2 mM Glutamax, 100 U mL^−1^ penicillin and 100 *µ*g mL^−1^ streptomycin and subsequently grown in this medium. Neurons were seeded in Poly-L-Lysine (PLL)-coated plates, using cells from two hippocampi (∼ 150, 000 cells) per 6 cm plate for protein and RNA extraction experiments. The day of dissection was counted as day *in vitro* 0 (DIV 0). At DIV 1, primary hippocampal neurons were infected with viruses expressing either RFP as control, or CRE-RFP to ablate UFM1 expression. Anisomycin (ANS) was obtained from SIGMA and dissolved in DMSO and flash frozen. Neurons were treated with 20 *µ*M ANS or DMSO for 2 h. All other reagents were obtained from Gibco or Sigma-Aldrich, unless stated otherwise.

### Immunoblotting analysis of mouse neurons

Cells were washed with PBS, lysed in RIPA buffer (150 mM NaCl, 1% (v/v) Triton X-100, 0.1% (v/v) SDS, 20 mM Tris pH 7.4–7.7) with protease inhibitors (1 *µ*g mL^−1^ aprotinin, 0.5 *µ*g mL^−1^ leupeptin, 17.4 *µ*g mL^−1^ PMSF) and 20 mM N-Ethylmaleimide (NEM) to prevent de-UFMylation. Protein concentration was determined using the Bradford method (Biorad). Proteins were then heated in a sample buffer at 65 °C for 20 min.

SDS-PAGE was performed with standard discontinuous gels or with commercially available 4%–12% Bis-Tris gradient gels (Invitrogen). Blot transfers were done using nitrocellulose membranes, according to standard procedures. Reversible MemCode protein staining (ThermoFisher) was performed to determine the total protein amount. Blots were blocked with a blocking buffer (PBS containing 5% (w/v) milk and 1% (v/v) Tween) and probed using primary and secondary antibodies diluted in blocking buffer. Blots were developed using enhanced chemiluminescence (GE Healthcare). Chemoluminescent signals were detected using the INTAS ECL Chemostar Imager. MemCode protein stain and blot images were analyzed using Fiji.

### RNA extraction and library preparation

Cytosolic and nuclear fractions from *Arabidopsis* and human RKO cells were homogenized and stored in TRIzol at −80 °C until further processing. RNA was extracted using the Direct-zol RNA Microprep Kit (Zymo Research) according to the manufacturer’s instructions. Mouse neuronal cell samples stored in 300 µL of DNA/RNA Shield were also kept at −80 °C until use. RNA from total cells of these samples was extracted using the Quick-RNA Miniprep Plus Kit (Zymo Research), following the recommended protocol. DNase I (NEB) treatment was performed during RNA isolation. Polyadenylated mRNA was enriched from total RNA using oligo(dT) magnetic beads (NEBNext Poly(A) mRNA Magnetic Isolation Module), followed by strand-specific library preparation using the NEBNext Ultra II Directional RNA Library Prep Kit for Illumina. All steps, including mRNA capture, first– and second-strand synthesis, end repair, adaptor ligation, and library amplification, were performed according to the manufacturer’s instructions. Library quality and fragment size distribution were assessed using an Agilent Fragment Analyzer. Final libraries were sequenced in single-end or paired-end 150 bp mode on an Illumina NovaSeq X Plus system at Vienna Biocentre NGS facility.

### RNA-seq processing and splicing analysis

Raw FASTQ files were preprocessed using Trim Galore with default settings to remove adapter sequences and low-quality bases. Trimmed reads were aligned using STAR^70^ in two-pass mode to improve splice junction detection. For *Arabidopsis*, alignments were performed against a genome index generated using the TAIR10 primary assembly and the Ensembl Plants Release 56 GTF annotation. For human RKO, reads were aligned to the GRCh38 primary assembly using the GENCODE v45 annotation. For mouse, reads were aligned to the GRCm39 primary assembly using the Ensembl Release 112 annotation. Transcript-level quantification was performed using Salmon with strand-specific settings, using the aligned BAM files as input^71^.

Aligned BAM files were sorted and indexed using samtools^72^, and splice junction files from STAR were used as input for rMATS^73^ to identify differential alternative splicing events between control and anisomycin-treated samples. rMATS was run in paired-end mode for *Arabidopsis* and human samples and in single-end mode for mouse samples, with strand-specific orientation and the variable read length option enabled. Splicing events were quantified across five canonical types: skipped exons (SE), alternative 5^′^ and 3^′^ splice sites (A5SS and A3SS), mutually exclusive exons (MXE), and retained introns (IR). Events were considered significant if they exhibited a false discovery rate (FDR) *<* 0.05 relative to wild-type and a difference in percent splice inclusion (|ΔPSI|) of at least 20% for SE events or 10% for all other event types. To define UFM1-dependent splicing changes, we selected events that were significantly altered in wild-type (FDR *<* 0.05) but not in the UFM1 mutant background (FDR *>* 0.05) under anisomycin treatment. For downstream analysis splicing events identified by rMATS were mapped to host Ensembl gene identifiers by overlapping event coordinates with a transcript database (TxDb) constructed from species-specific GTF annotations using the GenomicRanges and GenomicFeatures R packages^74^.

### Alternatively spliced transcripts characterization

To characterize the structural and functional features of alternatively spliced transcripts, genomic coordinates of splicing events were overlapped with transcript models derived from the reference GTF. Spliced regions were classified in a strand-aware manner relative to the coding sequence (CDS): segments upstream of the CDS start were defined as 5^′^ UTR, those overlapping the coding region as CDS, and those downstream of the CDS end as 3^′^ UTR. In cases of isoform ambiguity, a consensus classification was assigned using a priority hierarchy (CDS *>* 5^′^ UTR *>* 3^′^ UTR *>* non-coding *>* intronic). Subsequently, subcellular localization and presence of signal peptide were evaluated by querying Ensembl BiomaRt^75^. The presence of signal peptides (analyzed for human and mouse) was determined by annotated “SignalP” start positions, while subcellular localization was assessed using Gene Ontology (GO) terms for major compartments (e.g., Nucleus, Mitochondria, Endoplasmic Reticulum). Enrichment odds ratios for these features were calculated using Fisher’s exact test relative to the genomic background.

### Nonsense-mediated decay (NMD) susceptibility evaluation

To predict susceptibility to Nonsense-Mediated Decay (NMD), the genomic architecture of retained introns (IR) was analyzed for Exon Junction Complex (EJC) proximity and Premature Termination Codon (PTC) density. EJC deposition sites were inferred using splice junction density as a proxy from GTF annotations; genomic distances were calculated from the IR boundaries to the centers of all other introns within the same transcript. Downstream splice junctions located *>* 55 bp from the stop codon were classified as potential NMD-triggering EJC sites^41^. Additionally, the local sequence context of the 5^′^ and 3^′^ splice sites (±20 bp) was scanned for stop codon motifs (TAA, TAG, TGA) to quantify the density of potential PTCs introduced by the retention event.

### Splice site strength and motif analysis

To evaluate the strength of splice signals, the Maximum Entropy Model implemented in MaxEntScan^76^ was used to score 9-nt 5^′^ splice sites and 23-nt 3^′^ splice sites based on their adherence to position weight matrices derived from constitutive sites. Sequences flanking annotated junctions were extracted and processed using a Python wrapper. The ggseqlogo R package^77^ was used to visualize differences in consensus motif conservation between stalling-regulated retained introns and constitutive controls. Genomic sequences spanning the 5^′^ donor (20 bp exonic, 6 bp intronic) and 3^′^ acceptor (20 bp intronic, 6 bp exonic) junctions were extracted, with strand orientation computationally validated and corrected based on canonical GT-AG dinucleotides to ensure accurate alignment.

### Motif enrichment analysis

Motif Enrichment Analysis of RNA-binding proteins (RBP) and splicing regulatory elements was performed using AME (Analysis of Motif Enrichment) from the MEME Suite^78^, querying the CisBP-RNA database^79^. To establish a background control set, constitutive introns were extracted from human and mouse gene annotations. Sequences from UFM1-dependent and UFM1-independent splicing events were then individually tested for motif enrichment relative to this constitutive background. Analyses were restricted to specific genomic intervals: the 5^′^ splice site (spanning –20 exon/+6 intron nucleotides), the 3^′^ splice site (−20 intron/+6 exon nucleotides), and the remainder of the full intron body. Significant enrichment was determined based on E-values.

### Phylogenetic profiling of UFM1

Eukaryotic proteins families co-evolving with UFM1 were identified as described previously^80,81^. Firstly, we assembled a dataset of genome-predicted proteomes representing diverse eukaryotic taxa, as described previously^21^. The resulting proteins were compared to one another using Diamond BLASTP *v*2.1.11.165 (ultra-sensitive, maximum 250 hits, percent identity ≥ 25%, *E <* 10^−5^, query coverage ≥ 25%) and clustered based on similarity using MCL *v*22.282 using a variety of inflation parameters (*I* = 1.4, 2, 4, and 6)^82,83^. Initial clusters were established with an inflation value of 1.4, and clusters larger than 450 proteins were sub-divided based on higher inflation values when possible. To compensate for fragmented clusters, those with 20 assigned proteins or less were aligned using MAFFT *v*7.520 and profile hidden Markov Models (HMM) were generated^84,85^. These HMMs were used as queries to search the initial dataset using HMMER *v*3.4 (*E <* 10^−20^) and small clusters were merged with larger clusters (at least 50 sequences) when homology was detected, resulting in a final set of protein families.

To identify protein families which correlate with the presence of UFM1, the distribution of UFM1 was assigned across taxa based on previous curation^21^. To account for taxon sampling, species represented within the database were subdivided into supergroups, and monophyletic groupings of UFM1 presence and absence were defined. To calculate phylogenetic correlations, the median protein number for protein families with representation from at least 25 species was determined across UFM1 presence/absence clades and point-biserial correlations (PBC) were used to identify correlation coefficients between binarized UFM1 presence and median protein number^86^. Correlation significance was assessed using permutation tests (*n* = 100, 000) whereby protein families of equal size were generated through random sampling of the initial dataset without replacement. Given the independent generation of each test distribution, multiple test corrections were not applied. Correlating protein families were defined as those with a PBC ≥ 0.3 and *P <* 0.05. The distribution of correlated protein families was visualized using ITOL *v*6^87^.

To assess functional associations within UFM1-correlated protein families, all proteins included within the initial dataset were annotated with Gene Ontology (GO) terms. To this end, each protein was compared to the UniProt Knowledge-base using Diamond BLASTP (sensitive, maximum 100 hits, percent identity ≥ 25%, *E <* 10^−5^, query coverage ≥ 50%) and annotated using the Pfam database using HMMER (*E <* 10^−5^)^88,89^. Annotations were extracted from the resulting hits using Wei2Go^90^. GO-terms were assigned to individual protein families when at least 25% of proteins within a given cluster received the annotation. To identify GO-enrichments, term frequencies were compared between UFM1-correlated subsets and all eukaryotic protein families, and significance was assessed using permutation tests (*P <* 0.01*, n* = 100, 000).

### AlphaFold-Multimer Prediction

Protein-protein interactions between the UFMylation machinery and co-evolved candidates (*P <* 0.05, PBC ≥ 0.3) were predicted using AlphaFold-Multimer^91−93^. Predictions were executed using the ht-colabfold pipeline (https://gitlab.com/BrenneckeLab/ht-colabfold) on the Vienna BioCentre CLIP Batch Environment (CBE) cluster. Human protein sequences were retrieved from UniProt and supplied in FASTA format. Interaction confidence was assessed using the interface predicted TM-score (ipTM) and a custom PEAK score, calculated as the inverted and scaled (0 to 1) minimum Predicted Aligned Error (PAE) between chains, excluding intra-molecular interactions.

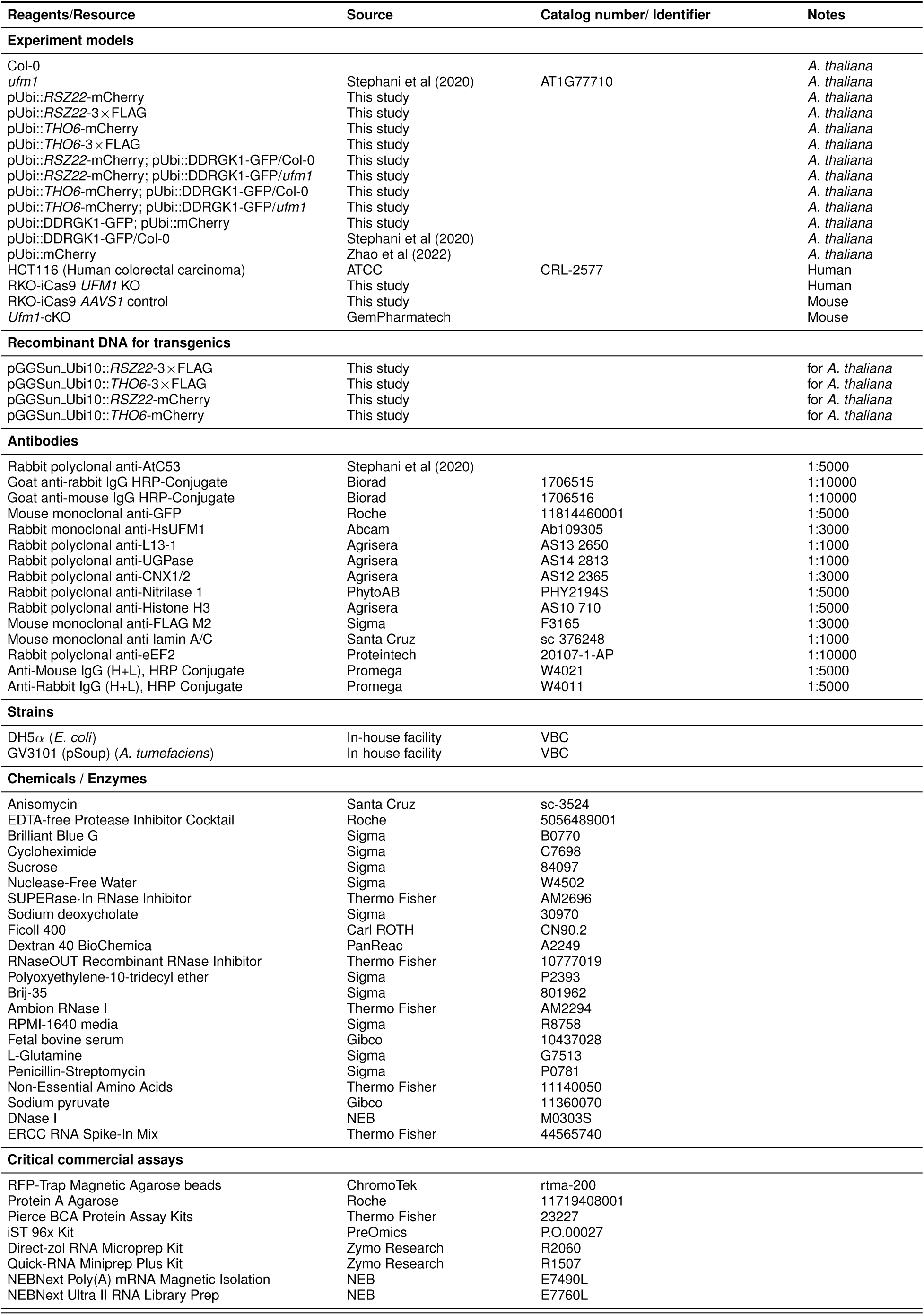
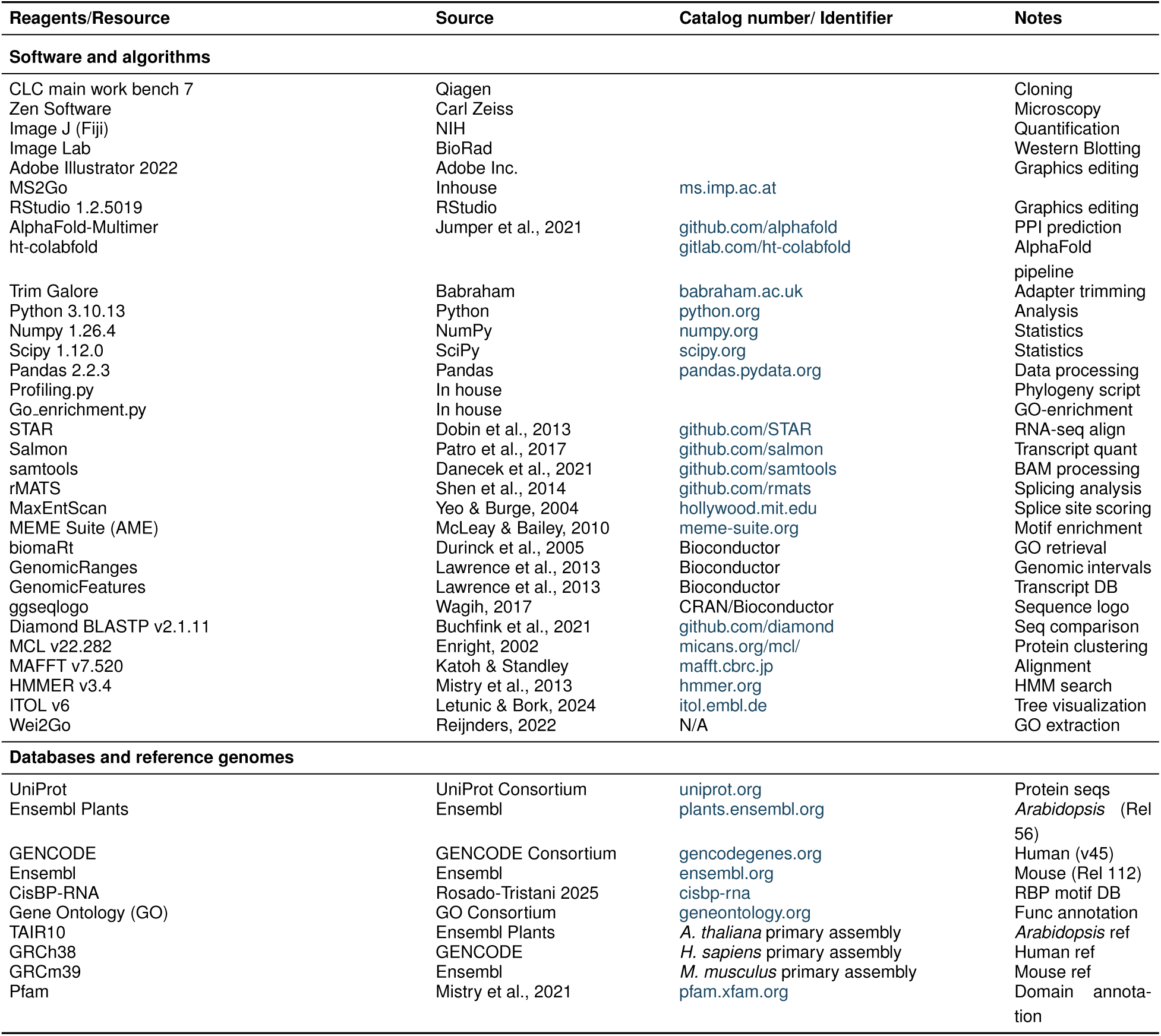
Key resources.

## Code availability

All custom code generated in this study for visualization, statistical analysis, and feature analysis is available at https://github.com/Papareddy/UFMylation_Ribosome_stalling_RNA_Splicing. Primary bioinformatics scripts were developed using human intelligence and subsequently streamlined via the Antigravity agentic framework (using Gemini or Claude Code) for redundancy mitigation.

## Data availability

All RNA sequencing data generated in this study are available at the National Center for Biotechnology Information Gene Expression Omnibus (NCBI GEO, https://www.ncbi.nlm.nih.gov/geo/) under accession number GSE325905. The mass spectrometry proteomics data have been deposited to the ProteomeXchange Consortium via the PRIDE^94^ partner repository with the dataset identifier PXD075540. All the raw data are available via Zenodo https://doi.org/10.5281/zenodo.19305729.

## Acknowledgements

We thank Vienna Biocenter Core Facilities (VBCF), particularly Proteomics, NGS, BioOptics and Plant Sciences. Proteomics analyses were performed by the Proteomics Facility at IMP/IMBA/GMI using the VBCF instrument pool. We acknowledge the use of high-performance computing resources provided by the bwHPC initiative and the Vienna CLIP computing cluster. The computational results reported within this publication were made possible by these infrastructures. We acknowledge funding from Austrian Academy of Sciences, Austrian Science Fund (FWF, P34944, I 6760, SFB F79), Vienna Science and Technology Fund (WWTF, LS21-009), European Research Council Grant (Project number: 101043370), DFG Heisenberg Program (Project number: 541762498) and Research Grants (Project numbers: 566190050, 568404408), the German research Foundation (DFG; SFB1286-A09 to N.B and M.T.). Ni Zhan was supported by a Marie Skłodowska-Curie Individual Fellowship (Project number: 101028611). We thank Adrian Söderholm (Versteeg lab) for his assistance with cytoplasmic and nuclear RNA fractionation. We are grateful to D. Warnecke, C. Harenberg, and F. Benseler for their expert technical assistance in mouse genotyping, and the staff of the Max Planck Institute for Multidisciplinary Sciences Transgenic Animal Facility for the generation and maintenance of mouse colonies. We also thank Sally Wenger for her technical support in confocal imaging. We thank Marion Clavel for helpful comments on the draft. We thank all members of the Dagdas lab for their support and helpful discussions for this project.

## Author contributions

N.Z., R.K.P., and Y.D. conceived and designed the project. N.Z. generated and characterized *Arabidopsis* transgenic lines, optimized the ribosome fractionation approach, and performed plant-based experiments including stress treatments, western blotting, co-immunoprecipitation, microsome, nuclei, and ribosome isolations. N.Z. prepared all samples for LC-MS/MS, conducted initial proteomics data analysis and target identification and validation, integrated datasets, and wrote the manuscript with input from all the authors. R.K.P. and H.A. established the plant microsome isolation pipeline and N.Z. continued to optimize it. R.K.P. optimized the *Arabidopsis* nuclei and cytosol isolation workflow. R.K.P. and N.Z. performed *Arabidopsis* nuclei and cytosol isolation for RNA sequencing. R.K.P. prepared RNA samples and libraries and led the RNA sequencing and analysis workflow. B.E. performed *Arabidopsis* root confocal imaging analysis and assisted with *Arabidopsis* transgenic line characterization and western blotting. A.S.A. and M.M. generated the human knockout lines, purified the cytoplasmic and nuclear fractions, and performed western blotting. A.S.A. prepared RNA from human samples for RNA sequencing. M.T. and C.P. performed mouse primary neuron experiments; N.B. supervised the mouse neural cell project. N.A.T.I. performed phylogenetic profiling and analysis. R.K.P. designed the rest of the software for data analysis and data visualization. R.K.P. and N.Z. assembled the figures. G.E.K. supervised the human cell work and provided input on the manuscript. Y.D. supervised the project, secured funding, and led the manuscript preparation and revision. All authors reviewed and edited the manuscript.

## Supplemental figures

**Fig. S1.**
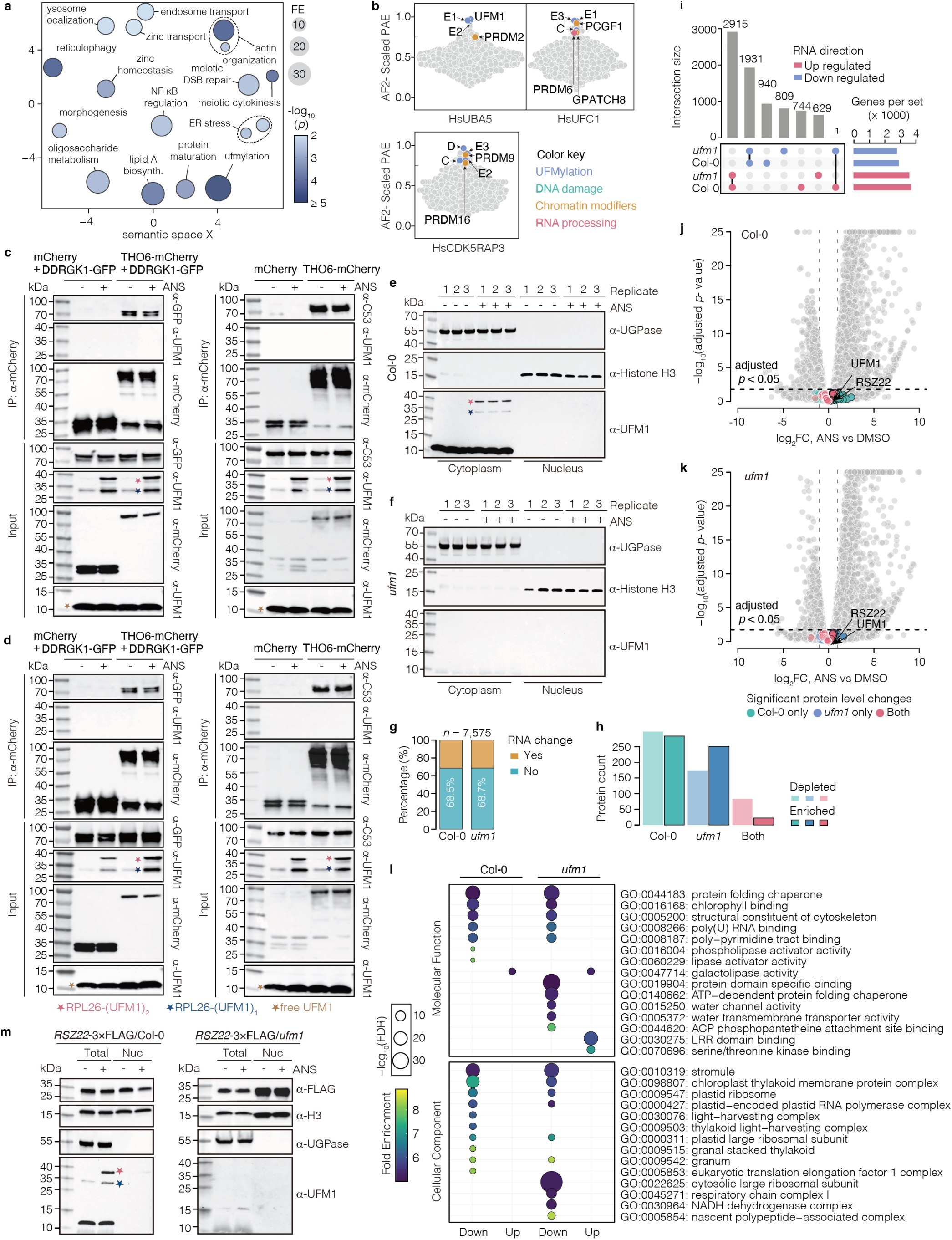
| Validation of UFM1 co-evolutionary candidates and transcriptomic-proteomic integration. **a**, Semantic clustering of GO terms for UFM1 co-evolutionary candidates (point-biserial correlation *>* 0.3, *P <* 0.05). Dot size and color indicate fold enrichment and *−* log_10_(*P*). **b,** Dot plot illustrating AlphaFold2 (AF2) Multimer interaction predictions of coevolved genes against UFMylation machinery. The Y-axis represents the average protein interaction score across the top three ranked models, calculated from domain-specific Predicted Aligned Error (PAE) values rescaled from 0 to 1. Abbreviations: P1, P2: UFSP1 and 2; E1, E2, E3: UBA5, UFC1, and UFL1; C: CDK5RAP3; D: DDRGK1. Coevolved genes with Scaled PAE *≥* 0.75 are colored according to key. **c,** *In vivo* co-immunoprecipitation (Co-IP) of THO6 interactions with DDRGK1-GFP and UFMylated RPL26 (left panel) or with endogenous C53 and UFMylated RPL26 (right panel) under DMSO or ANS (4 h) treatment. Proteins were extracted from the *Arabidopsis* lines co-expressing DDRGK1-GFP with either mCherry (control) or THO6-mCherry in Col-0 background seedlings (left panel), or from mCherry (control) and THO6-mCherry seedlings (right panel). Input samples were normalized by Bradford assay prior to loading. The consistent level of endogenous free UFM1 (unconjugated) in input fractions serves as an internal loading control to show equal protein loading across all lanes. THO6-mCherry was immunoprecipitated with RFP-Trap magnetic agarose, bound proteins detected by immunoblotting (*n* = 2). Mono-(1), di-(2) UFMylated RPL26 and free UFM1 are indicated. **d,** Biological replicate of (c). **e, f,** Immunoblotting of nuclear fractionation quality in 10-day-old Col-0 and *ufm1* seedlings (4 h ANS treatment). H3 and UGPase are nuclear and cytosolic markers, respectively. UFM1 blot confirms genotype-specific conjugates upon ANS as indicated (*n* = 3). **g,** Proportion of proteins showing RNA-dependent versus RNA-independent abundance changes in Col-0 and *ufm1* after ANS treatment based on integrated proteomic and RNA-seq analysis. **h,** Counts of proteins significantly enriched or depleted (*P <* 0.05) upon ANS treatment in indicated genotypes. **i,** UpSet plot showing intersections of nuclear DEGs (ANS versus DMSO) in Col-0 and *ufm1*. Bar heights indicate shared and genotype-specific upregulated (red) and downregulated (blue) gene set sizes, defining shared and genotype-specific transcriptional responses. **j, k,** Volcano plots of nuclear RNA-seq DEGs (ANS versus DMSO) in Col-0 (j) and *ufm1* (k). Genes with adjusted *P >* 0.05 are highlighted, indicating stable transcript levels and that many UFMylation-dependent proteomic changes are RNA-independent. **l,** GO enrichment of upregulated and downregulated DEGs in Col-0 and *ufm1*. Dot size and color denote *−* log_10_(FDR) and fold enrichment, respectively. Distinct transcriptional programs indicate that UFMylation dependent proteomic responses are primarily post-transcriptional. **m,** Immunoblot analysis of RSZ22-3*×*FLAG in total (whole-cell lysates) and nuclear fractions (replicate for Fig. 1i). 10-day-old RSZ22-3*×*FLAG seedlings (Col-0 and *ufm1* backgrounds) were treated with DMSO or ANS (4 h). H3 and UGPase serve as nuclear and cytosolic markers, respectively. Mono-(1) and di-(2) UFMylated RPL26 are indicated.

**Fig. S2.**
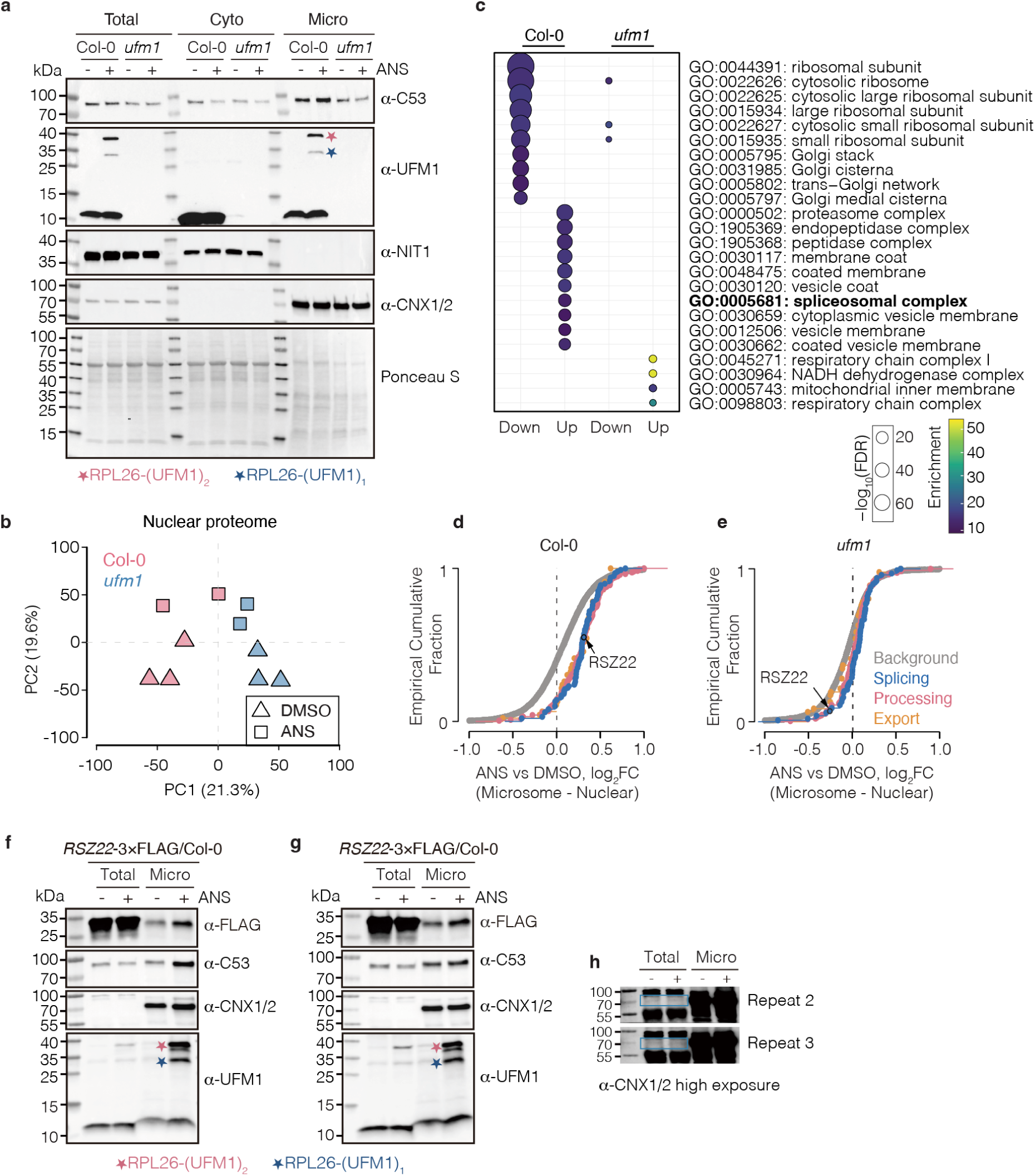
| Validation of microsomal enrichment and analysis of microsome-associated proteome. **a**, Validation of microsome isolation quality. Markers CNX1/2 (ER) and NIT1 (cytosol) confirm fraction purity. UFMylated RPL26 is enriched in the microsomal fraction. Mono-(1) and di-(2) UFMylated RPL26 are indicated. Ponceau S indicates total protein loading. **b,** PCA of nuclear proteomes (*n* = 3 for DMSO; *n* = 2 for ANS). PC1 and PC2 segregation reflects separation by genotype and treatment. **c,** GO enrichment analysis of proteins significantly altered in microsomal fraction upon ANS treatment in Col-0 and *ufm1*. Dot size and color denote *−* log_10_(FDR) and fold enrichment. **d, e,** Empirical cumulative distribution function (ECDF) plots showing subcellular redistribution. Δenrichment is defined as the difference in log_2_ fold change (ANS versus DMSO) between microsomal and nuclear fractions, for total proteome (grey) and RNA-related categories (colored) in Col-0 (d) and *ufm1* (e). RSZ22 is indicated as a representative example. **f–h,** Additional biological replicates for Fig. 2h. Immunoblot analysis of RSZ22-3*×*FLAG in total lysates and microsomal fractions. Mono-(1) and di-(2) UFMylated RPL26 are indicated. **h,** High exposure immunoblots of CNX1/2. *n* = 3 total biological replicates used for Fig. 2j quantification.

**Fig. S3.**
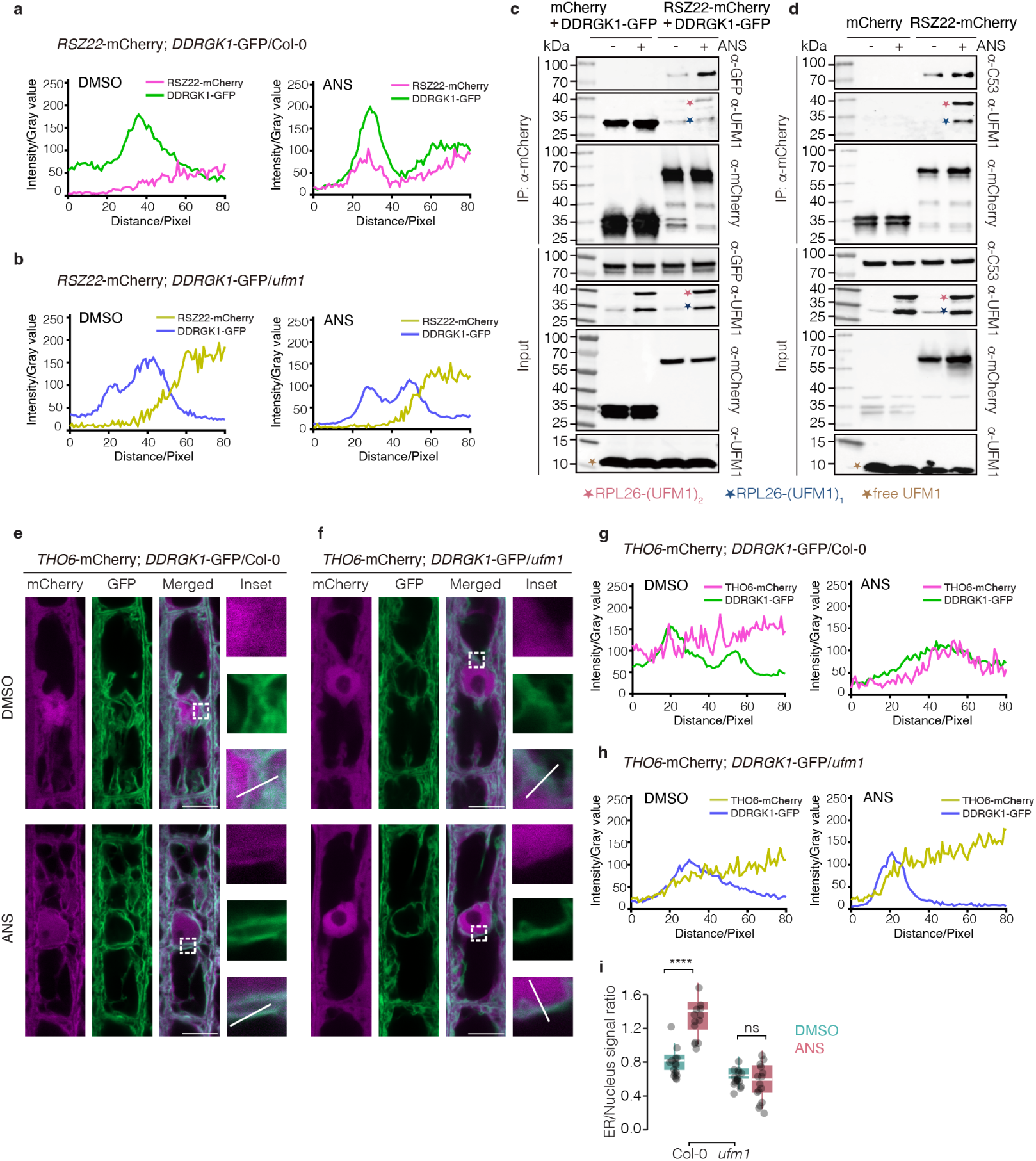
| UFMylation-dependent subcellular redistribution of RSZ22 and THO6, and RSZ22-UFMylation interac-tions. **a, b**, Fluorescence intensity profiles (line-scans) of RSZ22-mCherry and DDRGK1-GFP along the regions indi-cated in Fig. 2k, l. In Col-0 background (a), RSZ22 and DDRGK1 signals are shown in magenta and green, respectively, and ANS induces synchronized shifts and overlapping peaks. In *ufm1* background (b), RSZ22 and DDRGK1 signals are shown in yellow-green and purple, respectively, and the two signals remain spatially distinct regardless of ANS treatment. **c, d,** Representative biological replicates for Fig. 2n, o. *In vivo* Co-IP analysis of RSZ22 interactions with DDRGK1 and UFMylated RPL26 (c), or with endogenous C53 and UFMylated RPL26 (d), under DMSO or ANS (4 h) treatment. Lysates were incubated with RFP-Trap magnetic agarose, input and bound proteins were analyzed by immunoblotting. Mono-(1), di-(2) UFMylated RPL26 and free UFM1 are indicated. **e, f,** Confocal microscopy of THO6–mCherry and DDRGK1–GFP (ER marker) in roots from Col-0 (e) and *ufm1* (f) backgrounds. Five-day-old seedlings were treated with DMSO or ANS (1–2 h). ANS induces UFM1-dependent spatial overlap between THO6 and the ER in Col-0 but not in *ufm1*. Enlarged views and white lines indicate line-scan regions. Scale bars, 10 *µ*m. **g, h,** Fluorescence intensity profiles along the regions indicated in (e), (f). In Col-0 background roots (g), THO6 and DDRGK1 signals show overlapping peaks after ANS treatment, indicating co-localization. In *ufm1* background (h), the two signals remain spatially separated after ANS treatment. **i,** Quantification of ER-to-nucleus THO6 signal ratios from e, f. Data are mean *±* SD from 3 biological replicates. Each dot represents an individual cell (*n* = 15 cells from 3 biological replicates). Statistical significance was determined by two-tailed unpaired *t*-test **** *P ≤* 0.0001; ns, not significant.

**Fig. S4.**
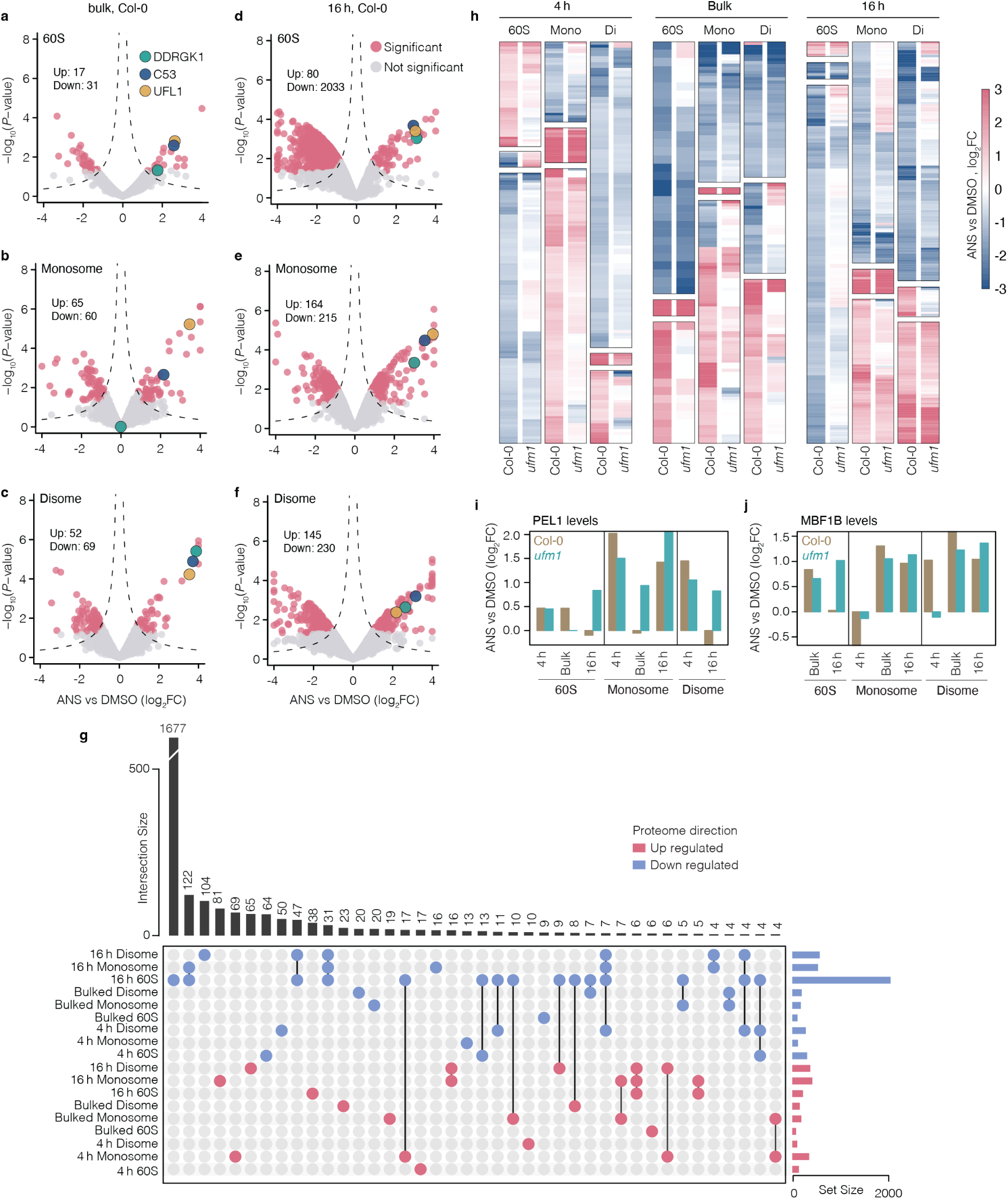
| Ribosomal proteomics evaluation and cytosolic RQC factor recruitment. **a–c**, Volcano plots from the bulk ribosome proteomics dataset (pooled from 1, 4, 8, 16 h treatments), showing ANS-induced recruitment of UFMylation components (UFL1, C53 and DDRGK1) to 60S (a), monosome (b), disome (c) fractions. x-axis, log_2_ fold change (ANS versus DMSO). y axis, *−* log_10_(*P* –value). **d**-**f,** Volcano plots from the 16 h ribosome proteomics dataset, showing recruitment of UFL1, C53, and DDRGK1 to 60S (d), mono-somes (e), and disomes (f). **g,** UpSet plot showing the intersections of significantly up-regulated (red) and down-regulated (blue) proteins across the indicated fractions (60S, Monosome, Disome) and conditions (Bulk, 4h, 16h). Vertical bars represent the size of the intersection; horizontal bars represent the total set size. **h,** Heatmap showing log_2_ fold changes (ANS versus DMSO) for proteins identified as significantly differentially abundant in Col-0 (WT). These same proteins are visualized in *ufm1* mutants to compare abundance patterns across time points (4h, Bulk, 16h) and ribosomal fractions (60S, Mono, Di). **i, j,** MS abundance profiles for PEL1 (i, plant homolog of mammalian PELO) and MBF1B (j, plant homolog of mammalian EDF1) across ribosomal fractions. Bars represent log_2_ fold change (ANS versus DMSO) in Col-0 (olive) and *ufm1* (cyan) from three datasets (4 h, bulk and 16 h). Both Col-0 and *ufm1* show comparable ribosomal recruitment of these two proteins.

**Fig. S5.**
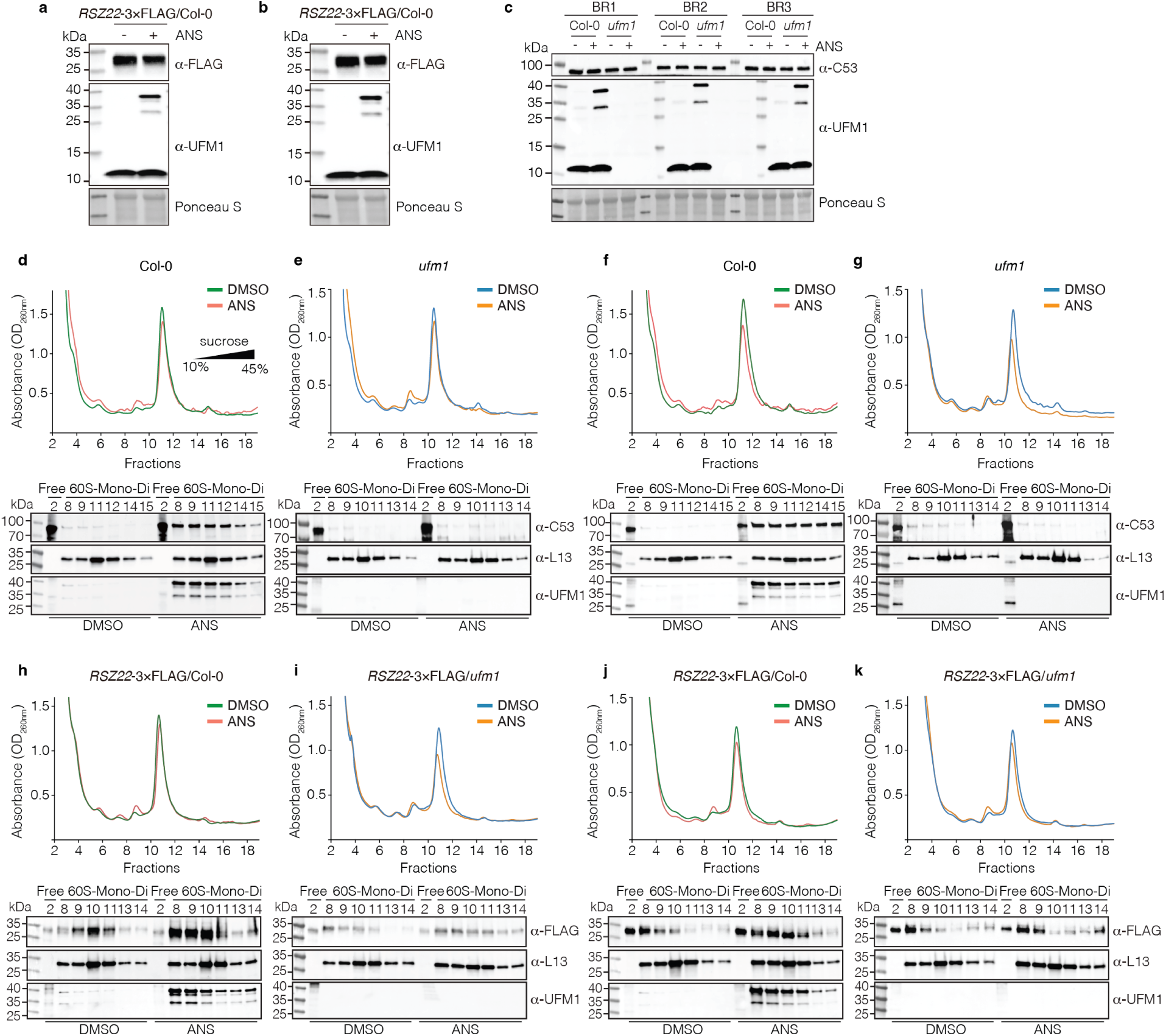
| Supporting evidence for the ribosomal recruitment of C53 and RSZ22. **a, b**, Immunoblot analysis of RSZ22-3*×*FLAG in total lysates of 7-day-old RSZ22-3*×*FLAG seedlings (Col-0 background) treated with DMSO or ANS (4 h). Lysates were probed using anti-FLAG and anti-UFM1 antibodies; Ponceau S indicates loading (*n* = 2 independent biological replicates). **c,** Immunoblot analysis of endogenous C53 in total lysates from Col-0 and *ufm1* seedlings treated with DMSO or ANS (4 h) (*n* = 3 independent biological replicates). **d–g,** Additional biological replicates of sucrose gradient sedimentation and immunoblot analysis of endogenous C53 in Col-0 (d), (f) and *ufm1* (e), (g). UV absorbance (A_260_) profiles (top) indicate ribosome distribution across 10%–45% gradients. Fractions were immunoblotted for C53, L13 (60S marker), and UFM1. Free: ribosome-free fractions. Results correspond to the second (d), (e) and third (f), (g) independent experiments, complementing Fig. 3f, g. **h–k,** Additional biological replicates of sucrose gradient sedimentation and immunoblot analysis of RSZ22-3*×*FLAG in Col-0 (h), (j) and *ufm1* (i), (k) backgrounds. UV absorbance (A_260_) profiles (top) indicate ribosome distribution. Fractions were immunoblotted for FLAG, L13, and UFM1. Results correspond to the second (h), (i) and third (j), (k) independent experiments, complementing Fig. 3h, i.

**Figure S6.**
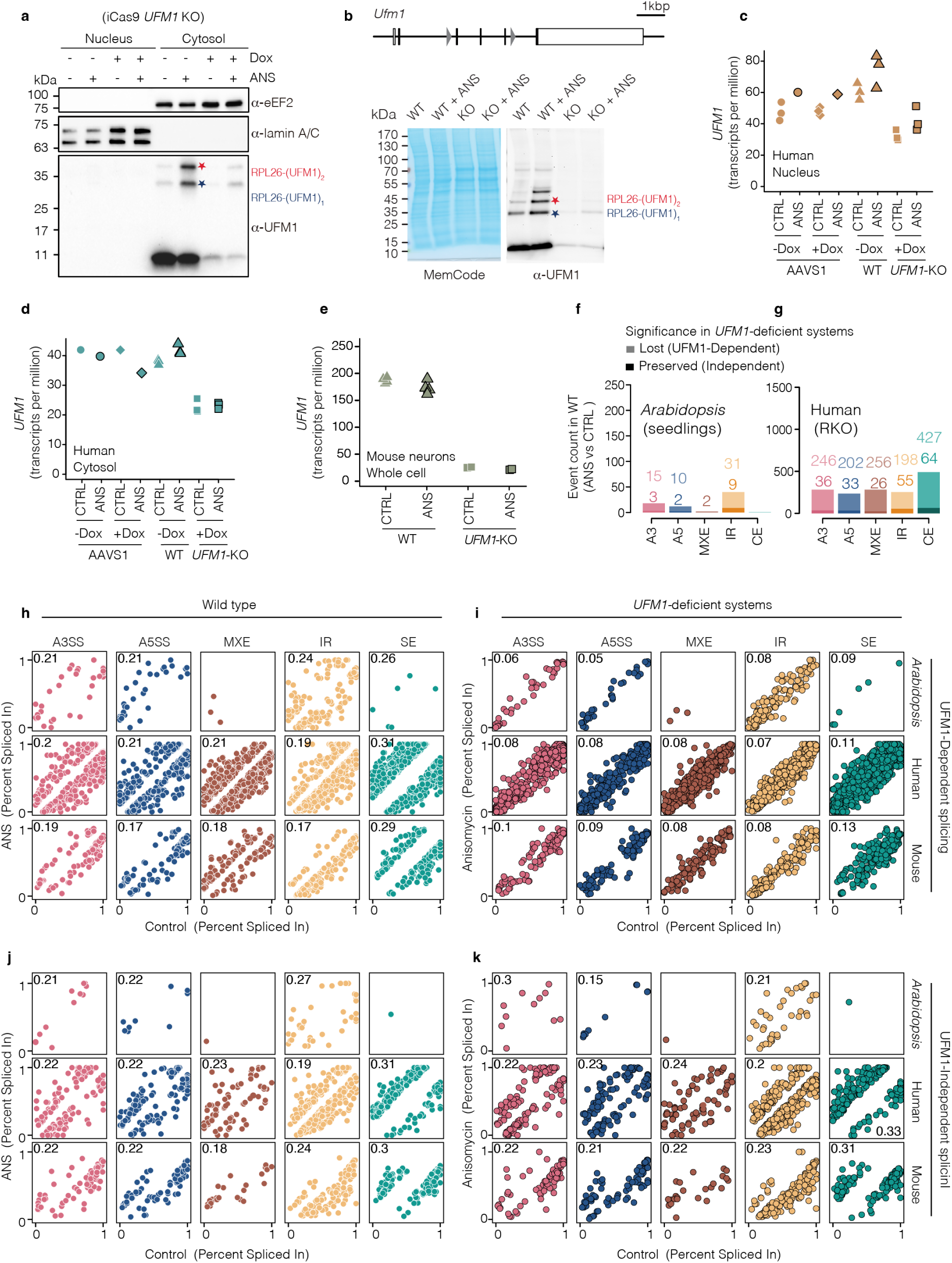
| Validation of subcellular fractionation and cross-species analysis of UFM1-dependent splicing. **a,** Immunoblot analysis of nuclear and cytosolic fractions in iCas9 *UFM1* KO cells treated as in Fig. 4b. Membranes were probed for eEF2, Lamin A/C, and UFM1. eEF2 and Lamin A/C serve as cytosolic and nuclear markers, respectively. Mono-(1) and di-(2) UFMylated RPL26 are indicated. **b,** Schematic and validation of the *Ufm1* conditional allele. (Top) Schematic representation of the *Ufm1* targeting strategy, indicating LoxP sites (grey triangles), coding exons (black boxes), and non-coding exons (white boxes); diagram is drawn to scale. (Bottom) Immunoblots (right) and total protein stain (left) of lysates from primary cortical neurons. Neurons were infected at DIV1 with lentiviruses expressing either RFP (WT) or Cre-RFP (KO) and treated at DIV12 with 20 *µ*M anisomycin (ANS) or DMSO vehicle for 2 h. Mono-(1) and di-(2) UFMylated RPL26 are indicated. **c, d,** *UFM1* transcript abundance (transcripts per million, TPM) in human nuclear (c) and cytosolic (d) fractions across AAVS1, WT, and *UFM1*-KO cell lines under control (CTRL) or ANS treatment conditions. **e,** *Ufm1* transcript abundance in whole cell lysates from mouse neurons comparing WT and *Ufm1*-KO under CTRL or ANS conditions. **f, g,** Quantification of ANS-induced alternative splicing events from the cytoplasmic fractions in WT *Arabidopsis* (f) and Human RKO cells (g), classified by event type. Lighter bars indicate UFM1-dependent events lost in UFM1-deficient systems; darker bars indicate UFM1-independent events retained. **h–k,** Scatter plots comparing PSI values between CTRL and ANS treatment for UFM1-dependent and independent events across *Arabidopsis*, human, and mouse transcriptomes. UFM1-dependent splicing events shown in WT backgrounds (h), and in UFM1-deficient systems (i) to demonstrate loss of splicing shift. UFM1-independent splicing events are shown in WT (j) and UFM1-deficient systems (k), demonstrating preserved splicing changes. Mean Absolute Error (MAE) values summarizing the global splicing shifts are indicated in each panel.

**Figure S7.**
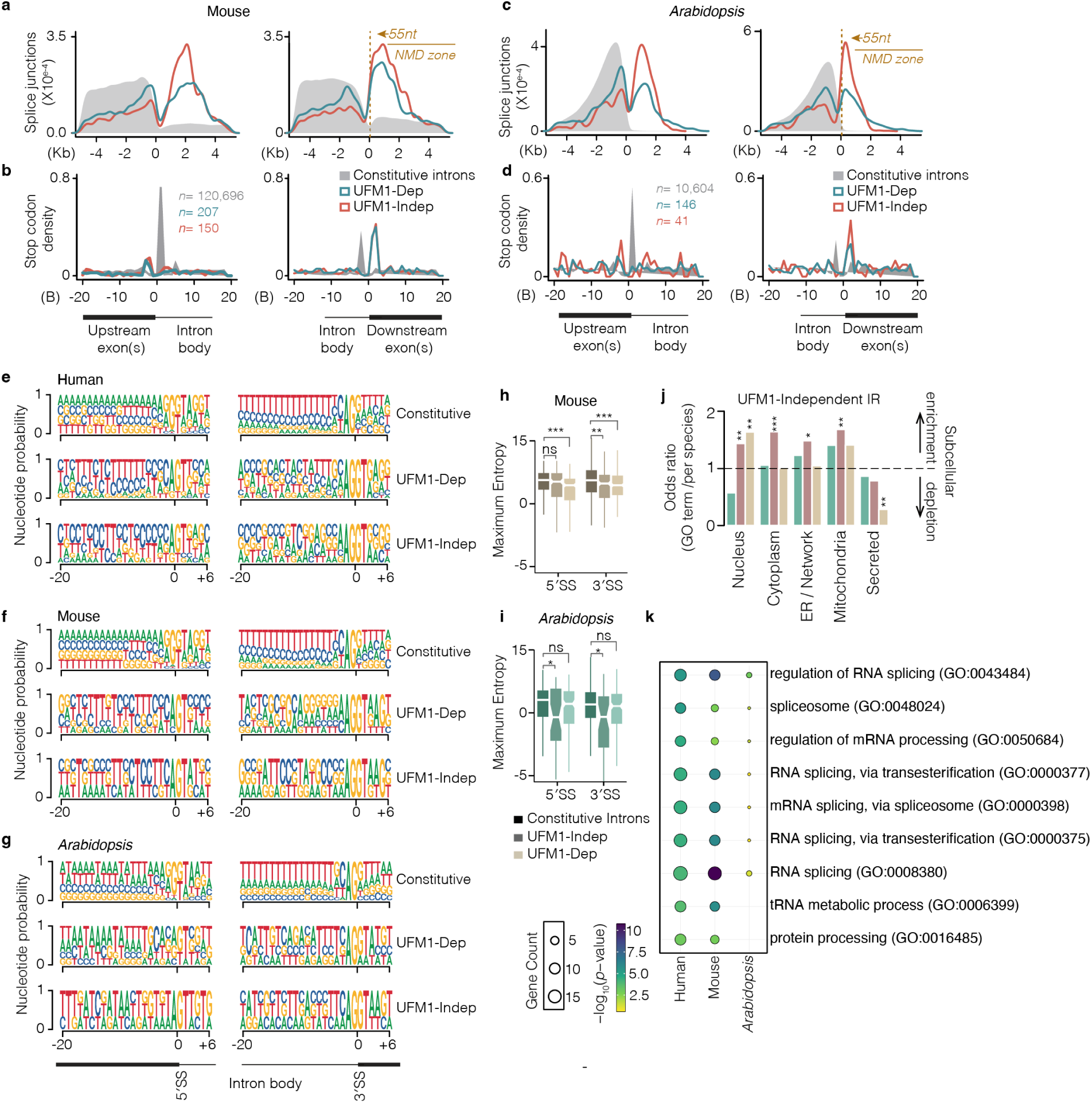
| Molecular features and functional enrichment of UFM1-associated splicing events. **a, c,** Metagene profiles showing the splice junction distribution relative to the stop codon in mouse (a) and *Arabidopsis* (c). Constitutive introns are shown in grey, UFM1-dependent events in blue, and UFM1-independent events in red. The 55-nt NMD zone is indicated by dashed lines and arrows. **b, d,** Density of stop codons in upstream exons, intron bodies, and downstream exon regions for mouse (b) and *Arabidopsis* (d). Comparison includes constitutive introns (grey), UFM1-dependent events (blue), and UFM1-independent events (red). Sample sizes (*n*) are provided for each category. **e–g,** Sequence logos displaying nucleotide probabilities at the –20 to +6 positions for 5_′_SS and 3′SS. Profiles are shown for constitutive, UFM1-dependent, and UFM1-independent introns in human (e), mouse (f), and *Arabidopsis* (g). **h, i,** Maximum entropy (MaxEnt) scores for 5′SS and 3′SS in mouse (h) and *Arabidopsis* (i). Box plots compare constitutive (black), UFM1-independent (dark green/grey), and UFM1-dependent (beige) introns. Statistical significance: *** *P <* 0.001; ** *P <* 0.01; * *P <* 0.05; ns, not significant. **j,** Odds ratios for subcellular localization (nucleus, cytoplasm, ER/Network, mitochondria, secreted) associated with UFM1-independent retained introns across three species. **k,** GO enrichment analysis UFM1-independent target transcripts, highlighting RNA splicing and protein processing functions in human, mouse, and *Arabidopsis*. Dot size indicates gene count; color gradient indicates significance (*−* log_10_ *P* –value). **l,** Comparison of RNA-binding protein (RBP) motif enrichment in UFM1-dependent versus UFM1-independent introns relative to constitutive introns from mouse neurons. Dot size denotes motif enrichment in UFM1-dependent introns compared to UFM1-independent introns.

